# Structural basis for target-site selection in RNA-guided DNA transposition systems

**DOI:** 10.1101/2021.05.25.445634

**Authors:** Jung-Un Park, Amy Tsai, Eshan Mehrotra, Michael T. Petassi, Shan-Chi Hsieh, Ailong Ke, Joseph E. Peters, Elizabeth H. Kellogg

## Abstract

CRISPR-associated transposition systems allow guide RNA-directed integration of a single DNA insertion in one orientation at a fixed distance from a programmable target sequence. We define the mechanism explaining this process by characterizing the transposition regulator, TnsC, from a Type V-K CRISPR-transposase system using cryo-EM. Polymerization of ATP-bound TnsC helical filaments explains how polarity information is passed to the transposase. Our Cryo-EM structure of TniQ-TnsC reveals that TniQ caps the TnsC filament, establishing a universal mechanism for target information transfer in Tn7/Tn7-like elements. Transposase-driven disassembly establishes delivery of the element only to unused protospacers. Finally, structures with the transition state mimic, ADP·AlF_3_, reveals how TnsC transitions to define the fixed point of insertion. These mechanistic findings provide the underpinnings for engineering CRISPR-associated transposition systems for research and therapeutic applications.

**One Sentence Summary:** Cryo-EM studies reveals the role of the AAA+ regulator TnsC for target-site selection in CRISPR-associated transposition systems.

## Main Text

CRISPR-associated proteins (Cas) are macromolecular machines that provide bacteria and archaea with adaptive immunity against bacteriophages and other invasive genetic elements. The RNA-guided DNA nuclease activity of CRISPR-Cas systems has been repurposed (most notably in the case of CRISPR-Cas9) for programmable genomic editing by making precise double-strand breaks in DNA (DSB) complementary to the RNA guide(*1*). Although conventional CRISPR-Cas systems can generate double-strand breaks with high fidelity at chosen DNA sites, the actual insertion of new DNA is dependent upon inefficient processes like homology-directed repair or nonhomologous end-joining. Moreover, introduction of a DSB into the host genome is dangerous as it can lead to genome instability.

Transposons are DNA insertion machines whose action does not involve an exposed DSB. Excitingly there exist examples of transposons that are naturally programable for targeting: Tn7-like transposons that have co-opted type I (Cascade)(*2*) and type V (Cas12)(*3*) CRISPR-Cas systems on multiple independent occasions for guide RNA-directed transposition. These CRISPR-associated Tn7-like transposons have been shown to exhibit a single programmable DNA integration event at a precise distance and in a specific orientation with respect to the protospacer adjacent motif (PAM) site(*4–6*).

In both prototypic Tn7 and in the Tn7-like transposon relatives which encode CRISPR-Cas systems, the overall mechanism of transposase recruitment and insertion into a specific target site remains elusive, with little structural information to guide mechanistic studies. Despite remarkable diversity(*7*), every RNA-directed transposition system characterized to-date contains a CRISPR effector domain (Cas12k in this study), target-site selector machinery (TniQ/TnsC), and transposase (TnsB) (***Figure 1A***). Previous structural studies have focused on the I-F3 Cascade-TniQ target-DNA binding complex, expressed from the element found in *V. cholerae*(*4, 8*). While the Cascade-TniQ structure reveals the physical association between the CRISPR effector domain and TniQ, how target-DNA binding ultimately results in transposition remains mysterious.

**Figure 1.**
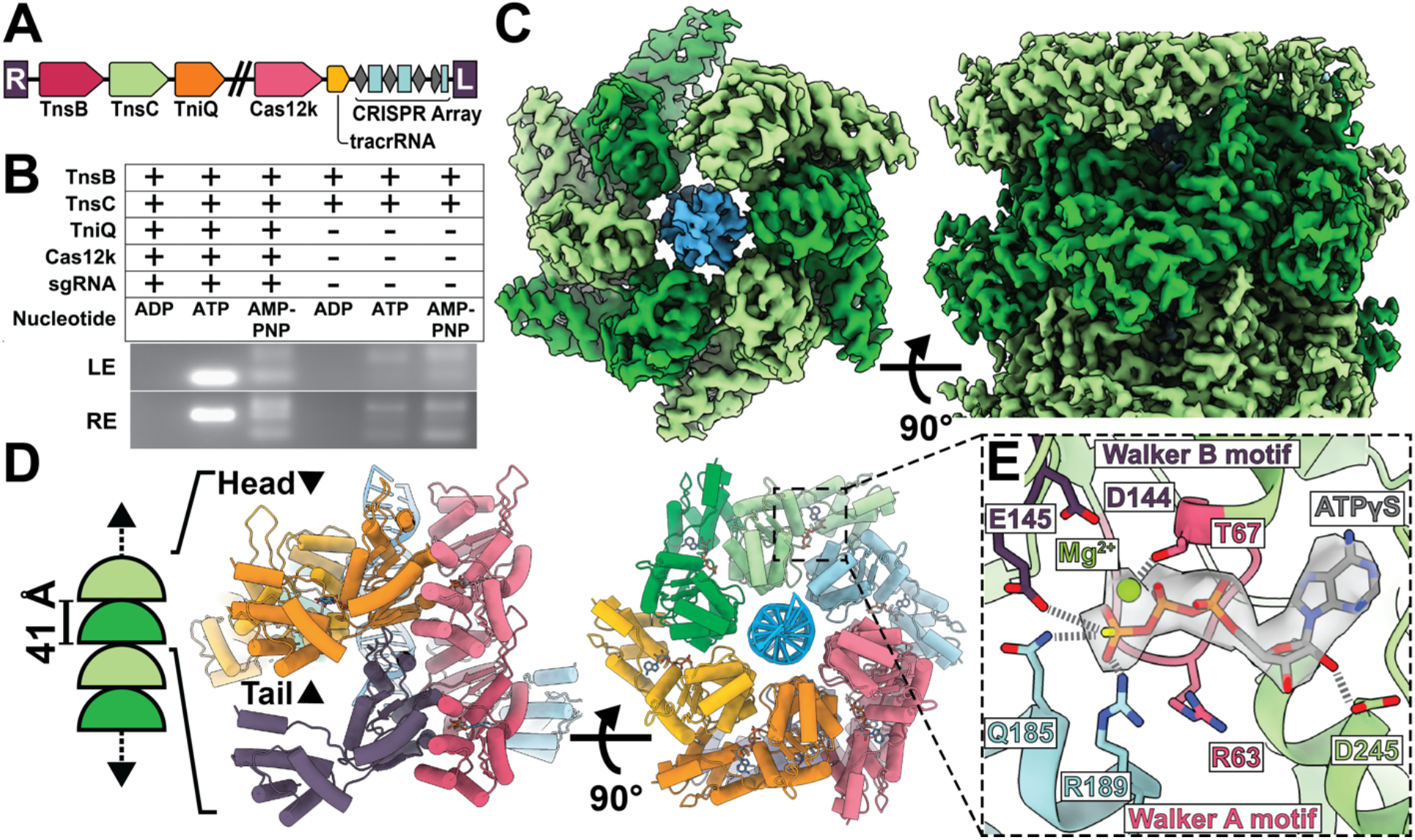
TnsC hydrolysis activity is crucial for ShCAST targeting and forms helical filaments in the presence of ATP. **(A)**. The ShCAST transposon contains: Cas12k RNA-binding module, TniQ, TnsC, and TnsB genes. **(B)** ATP-hydrolysis is required for targeted RNA-guided transposition. A single representative image of n=3 replicates is shown. (**C)** High-resolution (3.2 Å) cryo-EM reconstruction of ATPγS-TnsC forms a continuous helical filament encircling DNA. (**D)** Atomic model of TnsC helical filament consists of hexamers arranged in a helical spiral. The ‘head’ and ‘tail’ face of the hexamer are labeled. The arrangement of hexamers within the filament is referred to here as a ‘head-to-tail’ configuration (**E)** The ATP-binding pocket follows canonical features of AAA+ proteins, with conserved walker A (pink) and walker B motif (purple) coordinating ATPγS-binding. The adjacent subunit (light blue) forms inter-subunit contacts that contribute to coordination of the terminal phosphate.

CRISPR-associated transposons share crucial features with the prototypic Tn7 element. However, instead of a guide RNA complex, prototypic Tn7 uses the protein TnsD (TniQ domain containing protein) to recognize a specific attachment site (*attTn7*) in the bacterial genome for integration. An incompletely characterized interaction between TnsD and the regulator protein TnsC (a homolog of TnsC from RNA-directed transposition systems) will recruit the core TnsA + TnsB transposase bound to the ends of the element to integrate into the target DNA(*9*). TnsC is a AAA+ protein that has functional parallels with MuB, from bacteriophage Mu(*10*). More broadly, in both Tn7/Tn7-like systems and Mu, these AAA+ proteins are important regulators of transposition: mediating both (i) transposase recruitment to the target-site and (ii) preventing multiple insertions from occurring (also referred to as target-site immunity).

Here, we use cryo-EM to characterize TnsC of type V-K CRISPR-associated transposase system from S. *hofmanni* (referred to throughout as ShCAST). We were particularly drawn to the ShCAST system because of its relative simplicity (4 protein components), and established *in vitro* activity(*5*). We discovered that in the presence of ATP, TnsC forms a continuous spiraling helical filament bound to one DNA strand, providing a target site search mechanism that is also capable of conducting polarity information to the transposase. We demonstrate that propagation of the TnsC filament terminates when it forms a complex with TniQ, its binding partner, establishing a target site capture mechanism for Tn7 and the extended family of Tn7-like elements. We identify a TnsB transposase-directed process that disassembles the TnsC filament, driven by ATP hydrolysis, explaining how a targeted protospacer is only used once, while future insertions are diverted to new protospacers. This same TnsB interaction with TnsC would provide a mechanism to direct the TnsB-bound transposon DNA to a TnsC-TniQ complex for transposition. Notably, we find a post-hydrolysis TnsC mimic (i.e. ADP·AlF_3_ TnsC) collapses to a single hexamer that would be capable of conveying precise distance information from the protospacer to the point of insertion, and points to a nucleotide feature for stabilizing the TnsC-TniQ complex.

## Results

### ATP hydrolysis drives ShCAST target site selection

The TnsC regulator protein conveys essential information from the guide RNA complex to the transposase. Previous work with prototypic Tn7 and the Mu transposition system indicate that the nucleotide bound state of the TnsC and MuB ATPase is important for target site selection. Mu is a well-studied model system for transposition: MuB ATPase forms helical filaments in the presence of ATP, and MuB disassembly is stimulated by MuA transposase (*11, 12*). ATP hydrolysis is required for proper target selection in both Mu(*11*) and prototypic Tn7(*13*). To test whether ATP-hydrolysis is required for ShCAST targeting, we performed an *in vitro* transposition assay (See materials and methods)(*5*). Clearly targeted *in vitro* transposition was only detected in the presence of ATP (***Figure 1B***). Sequencing of seven independent events indicated transposition occurred at the expected distance from the PAM with two events being simple inserts and five co-integrates as found previously(*5, 14, 15*) (***Figure S1A***). Interestingly, a faint ladder of products could consistently be seen in the reaction mixture containing non-hydrolysable ATP-analog, AMPPNP (***Figure 1B***). This suggests a significant level of untargeted transposition could occur, a hypothesis we could confirm by analyzing the reaction products ***(Figure S1B-C)***. Robust targeted transposition required the presence of all reaction components, however we saw a faint ladder of PCR products in the TnsB and TnsC only lanes (***Figure 1B***). This is consistent with the untargeted products found in previous studies(*5, 16*). This indicates that, similar to Mu and prototypic Tn7, ATP-hydrolysis (via TnsC) is required for ShCAST to select the correct target(*17*). We also discovered that, similar to MuB ATPase(*18*), ShCAST TnsC forms helical filaments in the presence of ATP, AMPPNP, and ATPγS (***Figure 1C,S2-3***). To investigate the structural basis of these observations, we pursued the cryo-EM structure of TnsC.

### The atomic structure of TnsC possesses a canonical AAA+ fold and forms helical filaments

We discovered that TnsC is able to adopt a helical filament with a 6_1_ screw axis encircling DNA in the ATP (or ATPgS) bound states (***Figure 1C, S3***). Therefore, each TnsC hexamer has two potential polymerization interfaces, which we refer to as the ‘head’ and the ‘tail’ face (***Figure 1D***). Helical search of the cryo-EM images (using IHRSR) defines in a rise of 6.82 Å and a twist of 60°, consistent with helical layer-line analysis (***Figure S3B & S4A***). The ATPgS-bound state is of higher resolution overall (3.2 Å vs 3.6 Å for ATPgS vs ATP, respectively, ***Figure S5***) and with a more uniform distribution as assessed by local resolution estimates (3.0 - 4.0 Å, ATPgS vs 3.5 - 6 Å, ATP ***Figure S6***). A full-length model (257 of 276 residues) was built into the ATPgS cryo-EM map, starting from a homology model based on distantly related AAA+ YME1 (see Supp. Mat &Methods)(*19*) (***Figure 1D***).

ShCAST TnsC and prototypic Tn7 TnsC each have a conserved AAA+ domain, but ShCAST TnsC lacks the N- and C-terminal extensions of prototypic Tn7 which mediate its interactions with other Tns proteins(*20*) (***Figure S7***). As sequence analysis suggests, the structure follows most of the features of the initiator clade of AAA+ proteins. A DALI(*21*) search reveals strong structural resemblance to the N-terminal portion of Cdc6 (global rmsd is 2.7 Å)(*22*) (***Figure S8***), highlighting the high degree of conservation within the AAA+ domain, even among ATPases of highly divergent function. Correspondingly, the highly conserved Walker A motif (G^60^ESRTGKT^67^), and Walker B motif (M^140^LIIDE^145^) (***Figure 1E, S7***) form a pocket for the ATP binding. In addition to these intra-subunit contacts, the ATP-binding pocket is completed by R189 (the arginine finger) and Q185 from an adjacent subunit. These residues form hydrogen-bonding interactions with the terminal phosphate of ATP, both recognizing ATP and stabilizing helical filament assembly (***Figure 1E***). Notably, D144 appears too distant (4.6 Å) from the magnesium ion to facilitate a nucleophilic attack on the gamma-phosphate of ATP (***Figure 1E***), suggesting that a conformational change (possibly brought about by the transposase, TnsB) is required to carry out ATP hydrolysis that is relevant to recruitment of the TnsB bound ends and processes driving transpositions to new protospacers (see below and see discussion). These observations explain why TnsC, which is predominantly a monomer in solution, requires ATP to oligomerize into the observed helical filament, but can be readily disassembled upon ATP-hydrolysis.

### TnsC polymerization defines downstream transposition insertion polarity

ATP-bound TnsC forms a right-handed helical filament wrapping around the DNA duplex, forming a spiral ‘ladder’ of interactions with the sugar-phosphate backbone (***Figure 2A***). Each TnsC subunit contributes two amino-acid contacts: K103 and T121, to interact with two adjacent backbone phosphates (***Figure 2A***), which explains why ShCAST TnsC exhibits little to no DNA-sequence specificity, similar to MuB(*18*) and prototypic Tn7 TnsC. In the ShCAST system, these protein-DNA contacts distort duplex DNA to match the helical symmetry of the filament (***Figure 2B***). Strikingly, these interactions are formed preferentially with one strand of the DNA duplex (***Figure 2A***), and the local resolution of the complementary strand of DNA is substantially worse than the bound strand (***Figure S6B&D***). This suggests that TnsC will bind and assemble on single-stranded DNA similarly to double-stranded DNA, a hypothesis we confirmed using EM (***Figure S9***).

**Figure 2.**
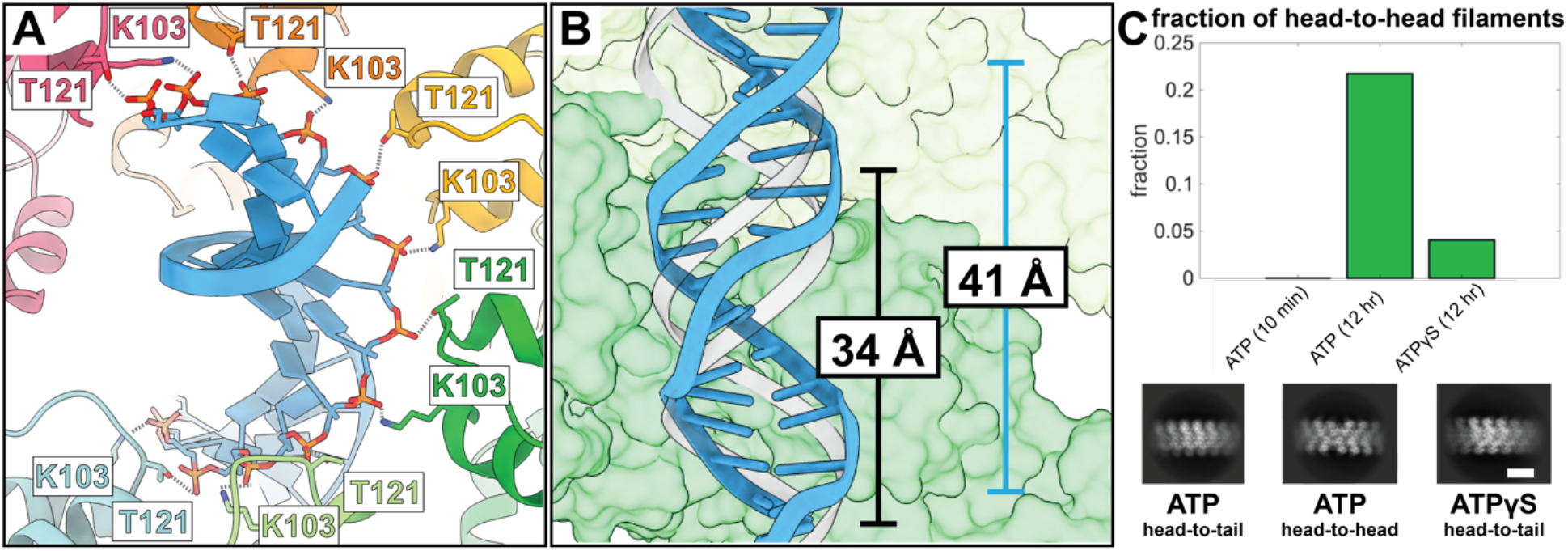
The TnsC filament searches for its target-site by selectively associating with one strand of the DNA duplex. **(A)** Each TnsC subunit makes two major contacts with the DNA sugar-phosphate backbone, forming a ladder of interactions selectively with one strand of DNA. (**B)** TnsC-DNA interactions result in DNA distortions. One full turn of B-form DNA spans 34 Å (gray), however the spacing between layers (41 Å) stretches the DNA, distorting the DNA to match TnsC’s helical spacing. (**C)** The formation of filament collisions, otherwise referred to as ‘head-to-head’ filaments, increases over time in the presence of ATP. The presence of such structures is influenced by both nucleotide (ATP vs ATPγS) and increase over time (more are present after 12 hours vs 10 min). Relevant 2D class averages for each sample are shown.

Filament polymerization is generally a unidirectional process. Consistent with this, TnsC filaments reconstituted with non-hydrolyzable analog ATPγS or frozen immediately upon reconstitution (with ATP) exhibit uniform polarity to cover the entire DNA substrate used in our analysis (i.e. each ‘head’ of TnsC interacts with the ‘tail’ of the adjacent TnsC hexamer, a.k.a. ‘head-to-tail’). While ATP-dependent filament were stable over short timeframes, they appeared to be more dynamic with prolonged incubation. For example, when samples are incubated overnight, we observe a substantial number of two converging filaments, forming head-to-head structures where they meet (20% vs none, for 12 hours and 10 min reconstitution respectively, ***Figure 2C&S10***). This is consistent with a dynamic process where the single filament we initially observed coating the entire DNA substrate could presumably partially disassociate allowing new converging filaments to form. Therefore, we hypothesize that polar growth of TnsC filaments in the 5’ to 3’ direction of the bound DNA strand is the searching mechanism that enables TnsC to search for its target site, which is defined by the TniQ-guide-RNA complex.

### TniQ interacts with TnsC to define the target-site

In prototypic Tn7 and Tn7-like systems, target information is conveyed from a TniQ domain family protein to TnsC(*23*). In most cases, the TniQ domain is fused to a DNA binding domain recognizing a specific DNA sequence(*7*). By contrast, in the RNA-guided transposition systems, the target site is chosen by a TniQ-associated guide RNA complex. In the case of I-F3 systems TniQ is positioned at the insertion site via its association with cascade(*8*). We propose here that TnsC filaments are perpetually searching for a target-site via a growing filament. An attractive model explaining how insertions can occur only on one side of an effector complex in a mechanism that sets orientation polarity, holds that collision with an appropriately positioned TniQ is central to the polarity conferring process. Therefore, diverse targeting mechanisms can be achieved by fusing TniQ to different DNA-binding domains, and in the case of guide RNA-directed systems, to RNA-binding CRISPR-effector domains. We believe this would serve as a unifying model accounting for diverse targeting mechanisms spanning both prototypic Tn7 and CRISPR-Cas systems. In order to explore this further, we reconstituted a simplified, minimal system to probe the possible role of TniQ to act as a target-site selection factor.

To directly visualize the interaction between TnsC and TniQ and their possible roles in target selection, we incubated TnsC and TniQ together in the presence of ATP and DNA and examined the resulting complexes by cryo-EM, generating a 3.9 Å resolution reconstruction (***Figure S4B&S5***). We find that TniQ selectively engages with the polymerizing face, capping the TnsC filaments (***Figure 3B***), consistent with the idea that the 5’ to 3’ directional propagation of the filament leads to productive interactions with the Cas12k-TniQ complex. We observe a total of two TniQ monomers; each copy interacts with two TnsCs (***Figure 3B***), even though theoretically three TniQ can be bound to the advancing TnsC filament. Each TniQ monomer also appears to be interacting with DNA, contacting the DNA strand that is not bound by TnsC (**Figure S11A**). Despite the high overall quality of the cryo-EM reconstruction, the local resolution of TniQ is too low for *de novo* model building (6-9 Å, ***Figure S11B***), possibly because of either sub-stoichiometric binding or mis-alignments brought about by the symmetrical nature of the TnsC filament. Nevertheless, homology models of TniQ’s functional domains (helix-turn-helix and zinc-finger or HTH and ZnF, respectively) each built from the I-F3 TniQ crystal structure and individually docked (PDB 6V9P) explain the cryo-EM density well (***Figure 3C***). Both the HTH and ZnF motifs appear to interact with the same region of TnsC. In the type I-F3 system TniQ associates as a homo-dimer, however in the ShCAST system TniQ is naturally found as essentially only the minimal TniQ domain, lacking a dimerization interface (***Figure 3A***). Correspondingly, we do not see substantial protein-protein interactions between the two copies of TniQ. However, reminiscent of the I-F3 system, the two copies of TniQ are oriented such that the N-terminus of one TniQ monomer is close to the C-terminus of the other (***Figure 3B***). The ATPγS TnsC atomic model explains the remaining cryo-EM density well, indicating that TniQ binding itself does not change TnsC helical parameters.

**Figure 3.**
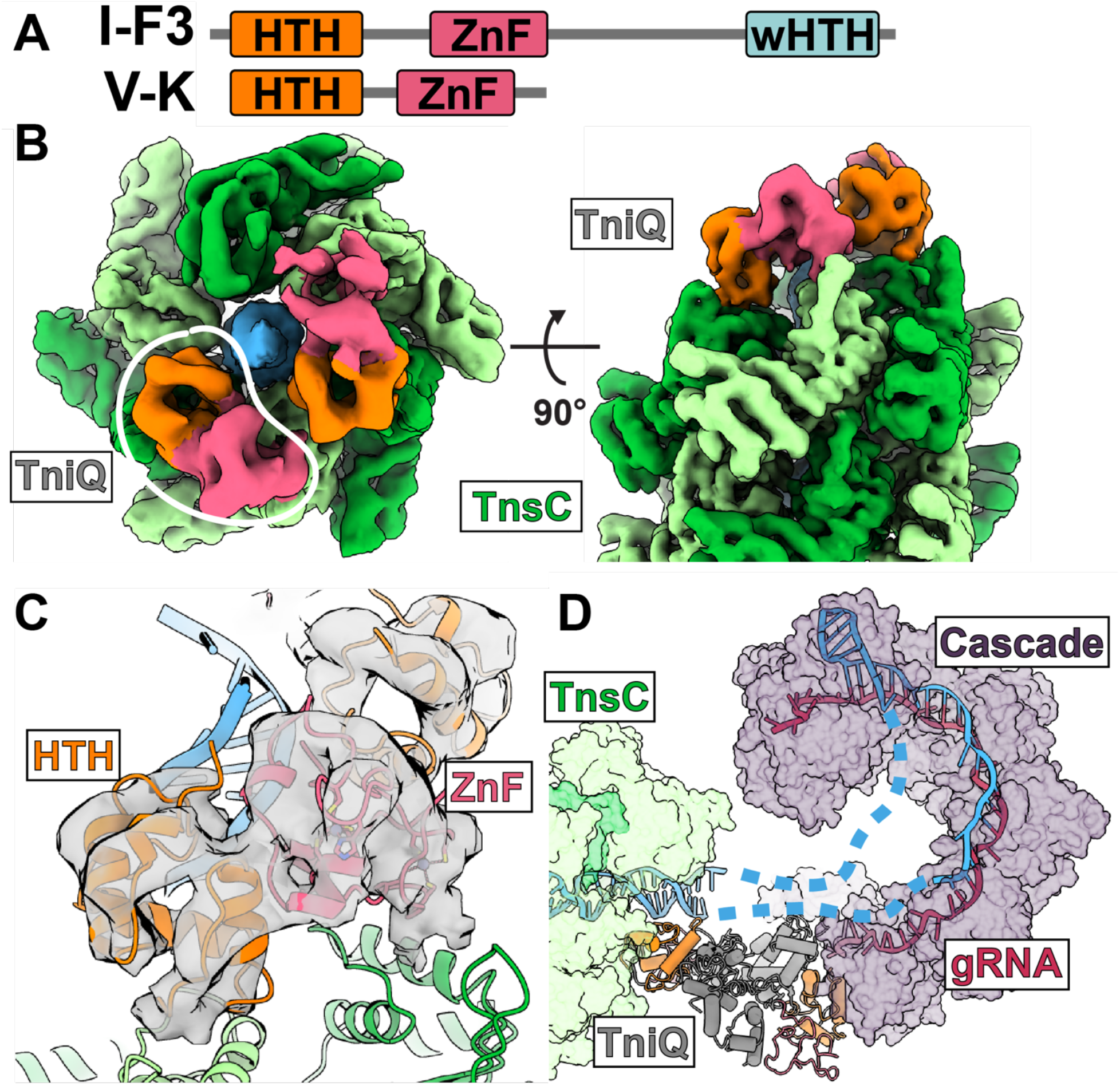
cryo-EM structure reveals how target-site selector protein TniQ conveys target-site information to polymerizing TnsC. **(A)** TniQ from type V-K CAST is truncated with respect to I-F3 TniQ. Functional domains corresponding to the helix-turn-helix (HTH, orange) motif and zinc-finger ribbon (ZnF, pink) motif are conserved. (**B)** Two copies of TniQ (orange/pink) interact with the head face of the ATP-bound TnsC filament. (right) Each monomer of TniQ interacts with two subunits (light/dark green) of TnsC. (**C)** Homology model of the helix-turn-helix (HTH) and Zinc-finger (ZnF) motif correspond well to the observed cryo-EM density map (**D)** Speculative higher order assembly of target-site binding complex, created by aligning Type IF-3 Cascade-TniQ (PDB: 6PIJ) with the TniQ-TnsC structure obtained in this study. This model reveals a pathway for PAM distal target-DNA binding.

The selective interaction of TniQ with only the advancing end of TnsC filaments explains how TniQ, likely also associated with Cas12k during the guide RNA-directed process, selects target-site insertion polarity. With this model of ShCAST TnsC-TniQ interaction in hand, we speculate on the possible higher order assembly of a guide RNA-directed target-site selection complex. Superimposing our docked ShCAST TniQ model onto the Type I-F3 Cascade-TniQ structure (PDB 6PIJ) reveals that the spatial organization of TniQ’s functional domains is conserved (global RMSD is 2.19 Å, ***Figure S11C***). Our model additionally reveals a clear pathway for the double-stranded DNA downstream of the R-loop (***Figure 3D***), which was not visualized in previous structures(*8*).

### The TnsC ADP**·**AlF_3_ structure represents a target-capture state and contains spacing information

A notable feature of Tn7 and Tn7-like elements, including guide RNA-directed systems, is that the point of element insertion is displaced a fixed distance from the actual machinery of target recognition, and no particular sequence is required for end joining on the target DNA. The TnsC filaments we identify here would provide a mechanism to offset the point of transposase association and point of insertion from the recognized target sequence. However, how can the precise spacing that is a hallmark of Tn7 and Tn7-like elements be dictated by an extended, continuous filament?

In prototypic Tn7 and Mu, TnsC (MuB) oligomers are disassembled by ATP-hydrolysis, stimulated by the transposase TnsB (MuA)(*12, 24*). We discovered that this feature is conserved in the ShCAST system: ATP bound TnsC filaments are disassembled upon addition of TnsB whereas AMPPNP bound filaments are not (**Figure S2**). This immediately suggested to us a link between ATP-hydrolysis and the precise insertional spacing downstream of the PAM site observed for all guide RNA-directed transposition systems to-date(*4, 5*). While a continuous TnsC filament would be incompatible with the insertional preferences observed (i.e. fixed spacing from the PAM site), TnsC filament ‘trimming’ by TnsB may result in a specific oligomeric configuration that neatly encodes spacing information. That is, the process that allows TniQ to stabilize the end of a filament could also stabilize a set number of terminal TnsC protomers from TnsB dissociation, instead channeling TnsB to form productive interactions for integration of the element.

To investigate the hydrolytic state, we determined the cryo-EM structure of TnsC using a nucleotide analog that represents a hydrolysis transition state mimic, ADP·AlF_3_. Our 3.8 Å cryo-EM reconstruction (***Figure 4A, S4C&S5***) revealed that ADP·AlF_3_ bound TnsC assembles only in an asymmetric structure that can be described as two hexamers oriented in a “head-to-head” configuration, similar to the configuration found when converging filaments meet (***Figure 2C,S10, S12A***). The structural configuration had obvious implications for relating the distance from the protospacer to the point of integration. Although the same length of DNA substrate was used for reconstitution of ADP·AlF_3_ and ATP-bound TnsC (60 bp in both cases), the ADP·AlF_3_ particles were significantly shorter (the DNA-binding footprint is 22 nucleotides total, ***Figure S12B&C***). This indicates that the ADP·AlF_3_ complex represents a conformational state of TnsC that is different from the continuous helical filament.

**Figure 4.**
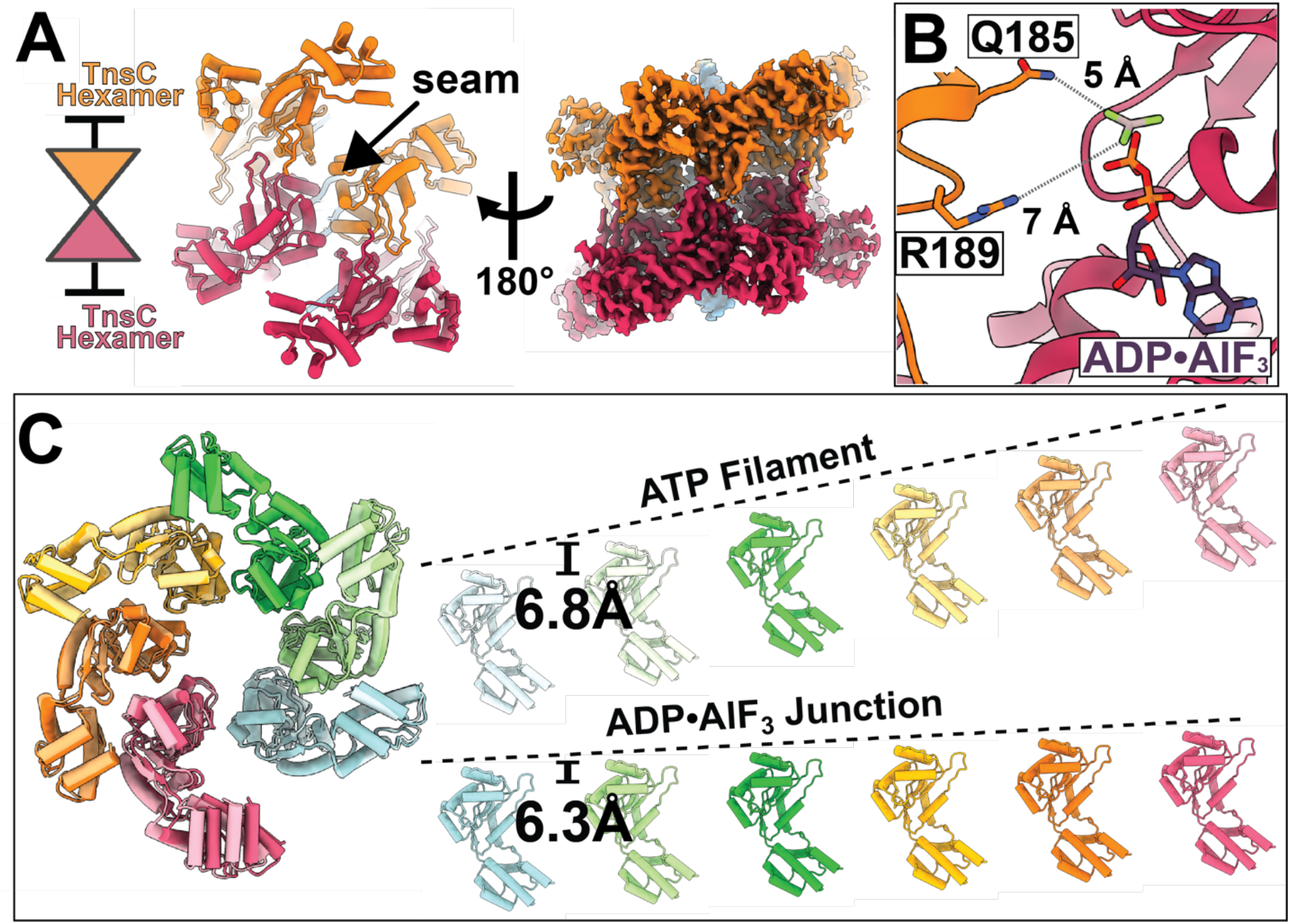
The post-hydrolysis state adopts a ‘closed’ structure that is incompatible with filament formation. **(A)** ADP-AlF3 3.7 Å cryo-EM density map reveals a head-to-head configuration of hexamers, in which the head face of one hexamer is oriented to interact with the other hexamer. This structure cannot support the formation of more than 1 helical turn. (**B)** TnsC subunits are repositioned in the ADP-AlF3 binding pocket such that the terminal phosphate is no longer coordinated by inter-subunit contacts. (**C)** The difference in subunit position results in a smaller rise (indicated by dotted lines) in the ADP-AlF3 junction (6.3 vs 6.8 Å per subunit) resulting in a ‘closed’ configuration that cannot accommodate another subunit to propagate the helical filament.

In the ATP-binding pocket of ADP·AlF_3_ structure, we do not see robust density corresponding to AlF_3_; the nucleotide density could alternatively be consistent with ADP·Mg (***Figure S13***). Nevertheless, the lack of a gamma phosphate results in a loss of inter-subunit contacts (Q185 and R189 are 5-7 Å from ADP), which results in an altered TnsC subunit organization (***Figure 4B***) and higher conformational flexibility (***Movie S1***). This altered binding-site configuration propagates to result in an overall smaller helical rise in the ADP·AlF_3_ state (6.3 Å vs 6.8 Å for ADP·AlF_3_ vs ATP, respectively ***Figure 4C***). We believe this represents the conformational changes that occur upon filament disassembly. The lack of TnsC filaments in the ADP·AlF_3_ sample also suggests that, upon ATP hydrolysis, the head-to-head configuration we observe is more stable against disassembly compared to the filament. This is intriguing because the interface between the two TnsC hexamers corresponds to the previously identified TniQ binding-site (***Figure 3B***). We speculate that the observed ‘head-to-head’ interface is substituting for the interface between TnsC and TniQ above and the Cas12k-TniQ complex found during bona-fide transposition. Thus, it is possible that one TnsC hexamer may remain stably bound to the Cas12k-TniQ complex after TnsB-stimulated ATP-hydrolysis.

## Discussion

Previous biochemical characterization of the ShCAST system(*5*) and work presented here indicates that programmed insertion of the DNA element occurs at a fixed distance downstream of the PAM. From these structural studies, we form a comprehensive picture that reconciles ShCast TnsC’s seemingly disparate proposed roles in target-site selection (***Figure 5***) which draws strong mechanistic parallels with Mu and Tn7. Our cryo-EM structures of ATP-bound TnsC reveals filaments that polymerize unidirectionally in the 5’ to 3’ direction. We hypothesize that such filaments represent a ‘searching’ state that would encounter the Cas12k-TniQ defined target-site with a specific polarity (i.e. downstream of the PAM). Our cryo-EM structure of TniQ-TnsC reveals the potential nature of this association at the target-site: only one face of TnsC forms productive interactions with TniQ. Importantly, we demonstrate that TnsB stimulates ATP-hydrolysis and filament disassembly which could draw TnsB (presumably bound to the transposon ends needed for integration) and to ‘follow’ TnsC to the chosen target-site. A similar process drives plasmid partitioning systems using ATPases(*25*). Interestingly, such a process could also serve to ‘chase away’ TnsC from the protospacer used for integrating to new protospacers. In the post-hydrolysis state, we observe that TnsC is incapable of forming a filament. While the exact form of TnsC in the active integration complex remains to be resolved, our results suggest how TnsC filaments interact with a Cas12k-TniQ complex with the right polarity and how TnsB-mediated ATP hydrolysis defines a shortened, integration-competent state.

**Figure 5.**
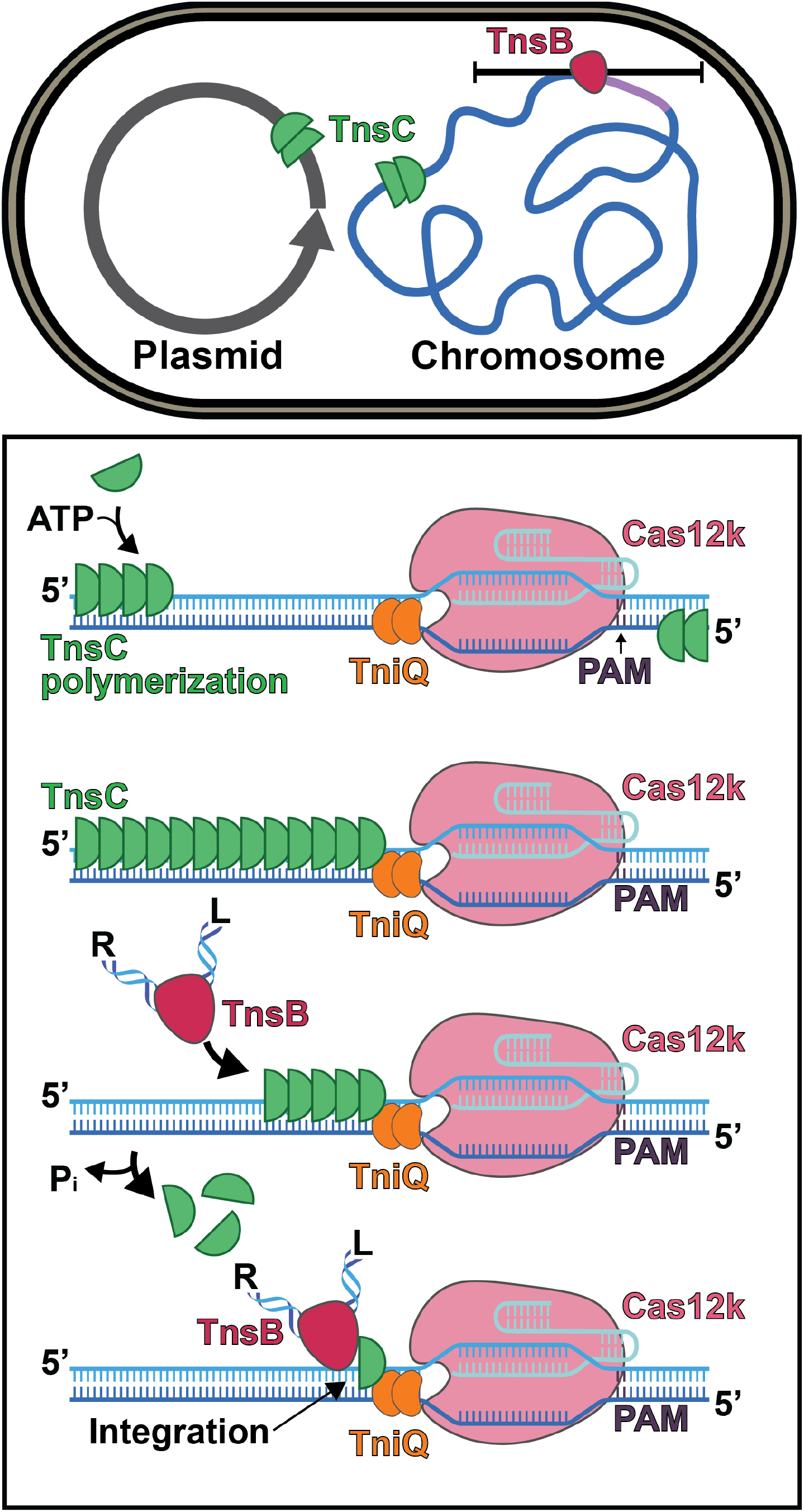
Mechanistic model describing TnsC’s role in target-site selection. TnsC promotes exploration of alternative target-sites by polymerizing along DNA. Previous insertion sites (indicated by TnsB binding) trigger TnsC depolymerization thereby rendering sites ‘immune’ to insertion. TnsC polymerizes unidirectionally along DNA on either strand in the presence of ATP. One direction will result in favorable interactions. Once TnsC encounters Cas12k-TniQ it is prevented from polymerizing further and forms a complex with TniQ. TnsB (bound to the terminal ends of the transposon) is then able to stimulate TnsC depolymerization and simultaneously be recruited to the target-site. The post-hydrolysis assembly of TnsC is disassembled to a finite oligomeric assembly (as opposed to a continuous helical filament), which allows integration a fixed distance from the protospacer.

We have also shown that TnsC induces a distortion in its DNA substrate by enforcing the helical parameters of the TnsC filament onto the DNA. Although the implications of this slight unwinding of the DNA require further exploration, it is tempting to speculate that this distortion could be crucial for its function. DNA distortions are a generally utilized driving force for integrases(*26*). As such, the high potential energy stored in the distorted DNA can be harnessed by the TnsB transposase machinery in order to facilitate forward transposition, or alternatively, may play a role in ensuring stable binding of TnsC at the target-site. In prototypic Tn7, it has been established that TnsABC transposes directly a specific distance in a single orientation of DNA distortions induced by TnsD(TniQ)(*27*). In the absence of TnsD, a DNA distortion associated with triplex DNA is sufficient to initiate TnsC recruitment and transposition(*28*). Our observation that TnsC necessarily introduces distortions in DNA upon binding to it might suggest that target-site selection complex (TnsD or Cas12k-TniQ) “pre-pays” the energy cost by distorting DNA such that TnsC can preferentially bind at the target site. Such a mechanism may also serve to stabilize TnsC binding post-hydrolysis.

This mechanism of filaments “searching” through DNA while being “chased” by the transposase bound to the ends of transposon provides an attractive, parsimonious model for explaining how a protospacer that is used for transposition can be subsequently ignored, and insertions driven to new protospacers for dispersal of the element(*6, 29*). While with canonical CRISPR-Cas systems the protospacer may be destroyed after being targeted by nucleases, insertions occur at a position offset from the protospacer in CRISPR-associated transposition systems, leaving it intact. The results presented here suggest that filamentation and disassembly of the AAA+ regulators in these systems is the structural basis of target immunity as described in other transposition systems with both Mu and prototypic Tn7. In particular, similarities observed here between the Mu and ShCAST systems indicates a conserved mechanism which might also be applicable to the prototypical Tn7 and I-F3 CRISPR-Cas-Tn7 systems. The physical association between TnsC and TniQ revealed by our cryo-EM reconstruction immediately explains how target-site selection information can be conveyed between the TnsC regulator and the RNA-binding CRISPR effector domain to result in programmable insertion. It is likely that analogous physical interactions occur in prototypic Tn7 and Type I-F3 guide RNA-directed systems.

Tn7-like transposons, including the prototypic Tn7 and guide RNA-directed transposition systems, display a remarkable diversity of targeting mechanisms which predominantly rely on proteins with TniQ-like domains. This diversity in targeting pathways suggests a remarkable adaptability for TniQ, which defines the target site for the core transposase(*3, 6, 29, 30*). In contrast, the core recombination components, TnsAB transposase and TnsC regulator, are likely to be highly conserved, both in structure and function. Notably, both TnsC and TniQ of Type V-K guide RNA-directed systems are significantly smaller than their equivalents from other systems (in prototypic Tn7 and other Tn7-like systems), containing only the highly conserved AAA+ core of TnsC and the HTH and ZnF motifs of the TniQ domain. Thus, we suspect that the structure visualized here likely represents the conserved, minimal functional interactions that are required between TniQ and TnsC. The interactions between ShCAST TnsB transpose and its regulator TnsC, however, remain to be determined. Therefore, the TnsC-TniQ machines whose structure we have defined here provide an excellent starting point for engineering new links and interactions between new target DNA recognition modules and the core transposase for more sophisticated genome-editing applications.

## Acknowledgements

We gratefully acknowledge the Cornell Center for Materials Research facility (CCMR), as well as Katherine Spoth and Mariena Silvestry-Ramos, for maintenance of electron microscopes used for this research (NSF-DMR1715879). We additionally acknowledge XSEDE for computational resources used for image processing (MCB200090 to E.H.K). We additionally thank Amanda Byer, Nozomi Ando, Gira Bhabha, and Seychelle Vos for advice on the use of nucleotide analogs, as well as members of the Ke and Peters group for helpful and stimulating discussions. We thank Lisa Eshun-Wilson, Eric Alani, and Brooks Crickard for valuable advice throughout this project. Last but not least, we thank Nancy Craig and Alba Guarné for valuable feedback on the manuscript.

## Author Contributions

E.H.K. conceived of the project and designed cryo-EM imaging experiments. J.P., and A.W.T. prepared samples for cryo-EM imaging. J.P. and E.M. analyzed, processed, and refined cryo-EM images to obtain 3D reconstructions. J.P., E.H.K., and E.M. built and refined atomic models. M.T.P designed and carried out the in vitro experiments under the guidance of J.E.P. All authors synthesized the ideas and contributed to figures and manuscript writing.

## Data and materials availability

Atomic models are available through the Protein Data Bank (PDB) with accession codes 7M99 (ATPγS TnsC), 7M9A (ADP·AlF_3_ TnsC consensus), 7M9C (ADP·AlF_3_ TnsC open), and 7M9B (ADP·AlF_3_ TnsC closed) ; all cryo-EM reconstructions are available through the EMDB with accession codes EMD-23724 & EMD-23725 (ATP TnsC), EMD-23720 (ATPγS TnsC), EMD-23721 EMD-23722 & EMD-23723 (ADP·AlF_3_ TnsC), and EMD-23726 (TniQ-TnsC).

## Supplementary Materials

### Materials and Methods

#### Protein Purification

Competent cells of *Escherichia coli* BL21-CodonPlus (DE3)-RIPL (Agilent) were transformed with the *tnsC*, the tnsB or the tniQ plasmid (Addgene). Purification of the proteins followed the same method. Single colonies were used to inoculate 10 mL LB media containing 34 µg·ml^−1^ chloramphenicol and 100 µg·ml^−1^ ampicillin and grown overnight at 37 °C as starter cultures. 1 L of 2xYT medium containing the same ratio of antibiotics were inoculated by the starter cultures. The cells were grown at 37 °C until mid-log phase before the temperature was lowered to 20 °C. The cells were then induced with 0.25 mM isopropyl-β-D-thiogalactopyranoside (IPTG) and continued to grow overnight (16-18 hrs) at 20 °C. Cells were then harvested by centrifugation at 5000 rpm for 10 minutes and resuspended in lysis buffer (50 mM Tris pH 7.4, 500 mM NaCl, 5% glycerol, 1 mM DTT, 1 mM PMSF, 0.1% NP-40 and 1 protease inhibitor/30 mL lysis buffer). Cell lysate was sonicated 7 times on pulse for 20 sec with 20 sec rest in between. The sonicated lysate was centrifuged for 30 min at 10,000 rpm at 4 °C. The supernatant was loaded onto a strep-tactin superflow resin column (iba) equilibrated with a loading buffer of 50 mM Tris pH 7.4, 500 mM NaCl, 5% glycerol. Bound protein was washed with the loading buffer with the addition of 1 mM DTT before the protein was eluted with the loading buffer containing 1 mM DTT and 5 mM *d*-desthiobiotin, with the exception of TniQ. Eluted TnsC or TnsB was then diluted to 200 mM NaCl, loaded on a heparin column equilibrated with 50 mM Tris pH 7.4, 200 mM NaCl, 1 mM DTT and 5% glycerol. The protein was step-eluted with 1 M NaCl. TniQ was eluted from the strep-tactin superflow resin with the SUMO tag removed by an overnight incubation at 4°C with SUMO protease at 1:100 weight ratio of protease to protein. TniQ was then concentrated and loaded onto a Superdex 200 Increase column (GE Healthcare) with a buffer of 25 mM Tris pH 7.4, 500 mM NaCl, 0.5 mM EDTA, 10% glycerol and 1 mM DTT.

#### *In vitro* Transposition Assay

Purified proteins were diluted to 2.5 µM in dilution buffer (500 mM NaCl, 50 mM Tris pH 7.4, 10% glycerol, 0.5 mM EDTA, 1 mM DTT); SUMO protease was added at 1/100 mass ratio to diluted SUMO-TnsC and incubated for three hours at 4°C. 2 mM ADP, ATP, or AMP-PNP were added to diluted and cleaved TnsC. sgRNA-PSP1 was prepared as described previously(*5*) and mixed with diluted Cas12k at a 2:1 sgRNA:Cas12k ratio (5 µM sgRNA). 100 uL reaction mixtures were prepared essentially as previously described(*5, 31*) with final concentrations of 0.54 nM pDonor_ShCAST_kanR (Addgene#127924),1.12 nM pTarget_CAST (Addgene#127926), 50 nM each TnsB, TnsC, TniQ, and Cas12K, 100 nM sgRNA-PSP1, 26 mM HEPES pH 7.5, 4 mM Tris pH 7.4, 40 mM NaCl, 10 mM KCl, 0.8% glycerol, 2 mM DTT, 50 µg/mL BSA, 0.04 mM EDTA, 0.2 mM MgCl_2_ and 2 mM ADP, ATP, or AMP-PNP as indicated. Separate donor (pDonor_ShCAST_kanR and TnsB) and target (pTarget_CAST, TnsC, TniQ, Cas12k, and sgRNA) reactions were prepared and preincubated at 37°C for 10 minutes before combining and adding MgOAc_2_ to a final concentration of 15 mM, then incubated for an additional two hours at 37°C. Reaction mixes were purified with GeneJET PCR purification kit and and assayed by PCR with primers JEP2486 + JEP2483 to monitor left end joining to target and JEP2482 + JEP2266 to monitor right end joining or by transforming cleaned up reaction mix into DH5α. Individual transformant insertion positions were determined by Sanger sequencing or PCR with JEP2486 + JEP2483 to detect left end joining as above.

JEP2486 : 5’ – CGTAGTATCTACGATACGTAGGAGG – 3’
JEP2483 : 5’ – CGCTGATGGGTCACGACGA – 3’
JEP2482 : 5’ – GCAAAGCGACAGCTAATTTGTCA – 3’
JEP2266 : 5’ – GAGGATGACGATGAGCGCATTG – 3’

#### TnsC filament reconstitution

Purified TnsC with the SUMO tag was dialyzed overnight at 4°C in the presence of SUMO protease (1/100 mass ratio) and 1/25 molar ratio of double-stranded DNA (a 60 base pair) against a reservoir of the following: 0.2-2 mM nucleotide (either ATP, ATPγS or ADP·AlF_3_), 200 mM NaCl, 2 mM MgCl2, 2% glycerol, 25mM HEPES, and 1 mM DTT. Sumo cleavage went to completion, as assessed using SDS PAGE. Then, the dialyzed mixture is concentrated and run over a sizing column (Superdex 200 Increase 10/300 GL, see Figure S1) or a spin-filter concentrator (amicon 50 kDa cutoff) with the following buffer: 25 mM HEPES, 200 mM NaCl, 2% glycerol, 1 mM DTT, 2 mM MgCl_2_, and 0.05 mM EDTA. For the fast reconstitution of the filaments, the sample was prepared within 10 minutes before the vitrification. The SUMO-tag removed TnsC was diluted to 30 µM into the buffer that contains double-stranded DNA substrate and ATP with the final concentration of 1.2 µM and 2 mM respectively. The composition of the dilution buffer was 25 mM HEPES, 200 mM NaCl, 2% glycerol, 1mM DTT, and 2 mM MgCl_2_.

#### TniQ-TnsC reconstitution

Purified TniQ was first buffer-exchanged with spin-filter concentrator (amicon 10 kDa cutoff) to the dilution buffer of following composition: 25 mM HEPES, 200 mM NaCl, 2% glycerol, 1mM DTT, and 2 mM MgCl_2_. Then the dilution buffer was supplemented with ATP and double-stranded DNA substrate to a final concentration of 2 mM and 0.6 µM, respectively. TnsC and TniQ were diluted into this buffer to the final concentration of 15 µM TnsC, 30 µM TniQ, followed by 4 hours of incubation in 37°C.

#### TnsB stimulated hydrolysis of TnsC

In order to assess whether TnsB stimulates ATP hydrolysis, we reconstituted ATP-bound or AMPPNP-bound TnsC filaments, as described in the previous section. Filaments were either incubated with a 1:1 molar ratio of TnsB or an equivalent volume of TnsB buffer. TnsB was purified according to previously described methods(*5*) except the sumo-tag was not removed. Incubation at 30° C for one hour was followed by negative stain EM. Briefly, 4 µL of sample was incubated on a continuous carbon grid(Electron Microscopy Sciences) before washing 3 times in a 50 µL droplet of milliQ water, next by embedding 3 times in 50 uL droplet of 2% Uranyl Acetate stain. Images were collected on a Morgagni (100 kV) at a magnification of 89,000x.

#### Cryo-EM sample preparation and imaging

reconstituted filament sample (ATP-bound, ATPγS-bound, and TniQ bound) was concentrated with the Amicon ultra-0.5 centrifugal filter unit (EMD Millipore) to 0.5 mg/mL. 3.5 uL of the concentrated sample was loaded onto the freshly plasma cleaned UltrAuFoil R1.2/1.3 holey gold grid (Quantifoil). ADP-AlF3 sample TniQ-bound TnsC filamentsSamples were vitrified using the Mark IV vitrobot (ThermoFisher) set to 100% humidity and 4°C settings. Samples were blotted for 6 seconds before being plunged into the liquid ethane cooled to −180°C using liquid nitrogen. The vitrified samples were imaged with a Talos Arctica (ThermoFisher) operating at 200 keV equipped with a Gatan K3 direct electron detector and Gatan bioquantum energy filter using a nominal magnification of 63,000X. The microscope was carefully aligned according to recommended procedures for coma-free alignment and parallel illumination settings(*32, 33*). 2910 and 3493 micrographs were acquired for ATP- and ATPɣS-bound filaments, respectively, with 3 by 3 image shift using the SerialEM(*34*) with the nominal defocus from −1.0 μm to −2.5 μm, and the total dose of 45 e^-^/Å^2^ (43 frames during 2.8 seconds exposure) for ATP-bound and 40 e^-^/Å^2^ (40 frames during 2.5 seconds exposure) for the ATPɣS-bound sample.

#### Image-processing, Model Building, and Model Validation

Beam-induced motion correction and the contrast transfer function (CTF) estimation of all the dose-fractionated micrographs were done using Warp(*35*). For the ATP-bound filaments, 2,910 micrographs were imported to Relion 3.1(*36*), and 2,700 particles were manually picked to generate a template for auto picking, which yielded initial 579,028 particles. The rise and the diameter of the filaments were obtained from the 2D classification using Relion 3.1. Simulated helix by relion_helix_toolbox was used as a reference in order to minimize the reference bias while using the known helical parameters. Initial alignment was done using the cryoSPARC homogeneous refinement(*37*), followed by 3D sorting in Relion 3.1(He and Scheres) to get the final stack of 33,312 particles. 3D variability analysis was carried out using Cryosparc(*38*). The combination of Relion3.1 CTF refinement, Bayesian polishing, and the helical refinement improved the resolution from 4.6 Å to 3.6 Å. For the ATPɣS dataset, 3,493 micrographs were motion-corrected and CTF-estimated with Warp, and the initial particle stack of 1,235,415 particles were picked using the pre-trained Warp BoxNet(*35*). Initial alignments from cryoSPARC were exported to Relion3.1 for further processing. Removing overlapping particles, and 3D classification with Relion3.1 resulted in 99,247 particles, which was subsequently improved by helical refinement, Bayesian polishing and CTF refinement to the final reported resolution (3.2 Å). For the ADP·AlF_3_ bound TnsC as well, motion-correction, CTF-estimation, and particle picking was done using Warp. Initial 754,099 particles were then subjected to 2D classification and ab-initio reconstruction in cryoSPARC, in order to generate the structure without reference biasing. Further processing in Relion 3.1 including the series of 3D classifications, CTF refinements, and Bayesian polishing resulted in the 3.8 Å resolution with the final 97,222 particles. For TniQ bound TnsC and fast-reconstituted TnsC filament dataset, essentially same pipeline was used.

The initial model was generated using the Phyre2 server(*39*), based on YME1 (PDB: 6AZ0). The initial model was docked into the cryo-EM density map and rebuilt manually using coot(*40*). Extended loops were rebuilt into the ATPɣS cryo-EM map using RosettaES(*41*). ATP and Magnesium atoms were docked in manually using UCSF Chimera(*42*). The full helical ensemble was subjected to symmetry constrained energy-minimization using Rosetta(*43*). DNA duplex was built in using coot(*40*) and refined into the cryo-EM map using phenix(*44*). TniQ model was built using the published crystal structure as a template. Helix-turn-helix (HTH) motif and Zinc-finger (ZnF) motif are built with I-TASSER server(*45*) and GalaxyWEB server(*46*) respectively. HTH motif was docked in manually followed by the real-space refinement of phenix into the cryo-EM map.

**Fig. S1.**
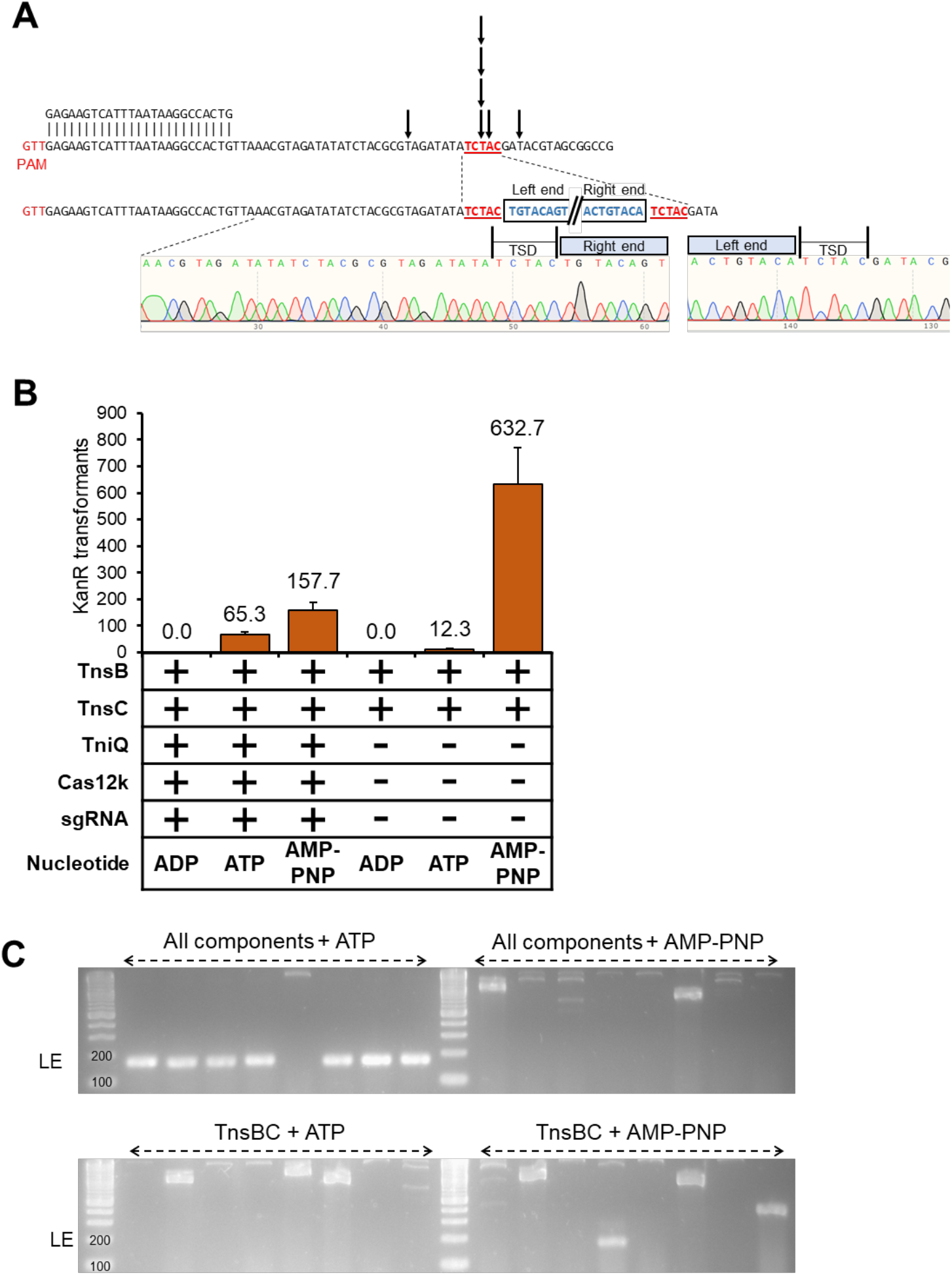
ShCAST RNA-guided transposition *in vitro* requires ATP. **A.** Sanger sequencing indicates seven individually isolated transposition events occur at the predicted distance from PAM (59 bp x4, 49 bp, 60 bp, 64 bp) with one representative trace shown. **B.** Colony counts from transformation of each transposition reaction are shown (one third of total reaction volume was used in transformation). Data indicate mean +/- standard deviation (n=3). **C.** Insertion position of eight individually isolated transposition events from the listed conditions was determined by PCR amplification with primers binding to the transposon left end and the expected target site in pTarget as in Figure 1B. An on-target insertion would produce a product size ∼168 bp; seven out of eight with all transposition components and ATP are predicted to be in the correct site and orientation while no insertions are predicted to be in the correct site in the other conditions assayed.

**Fig. S2.**
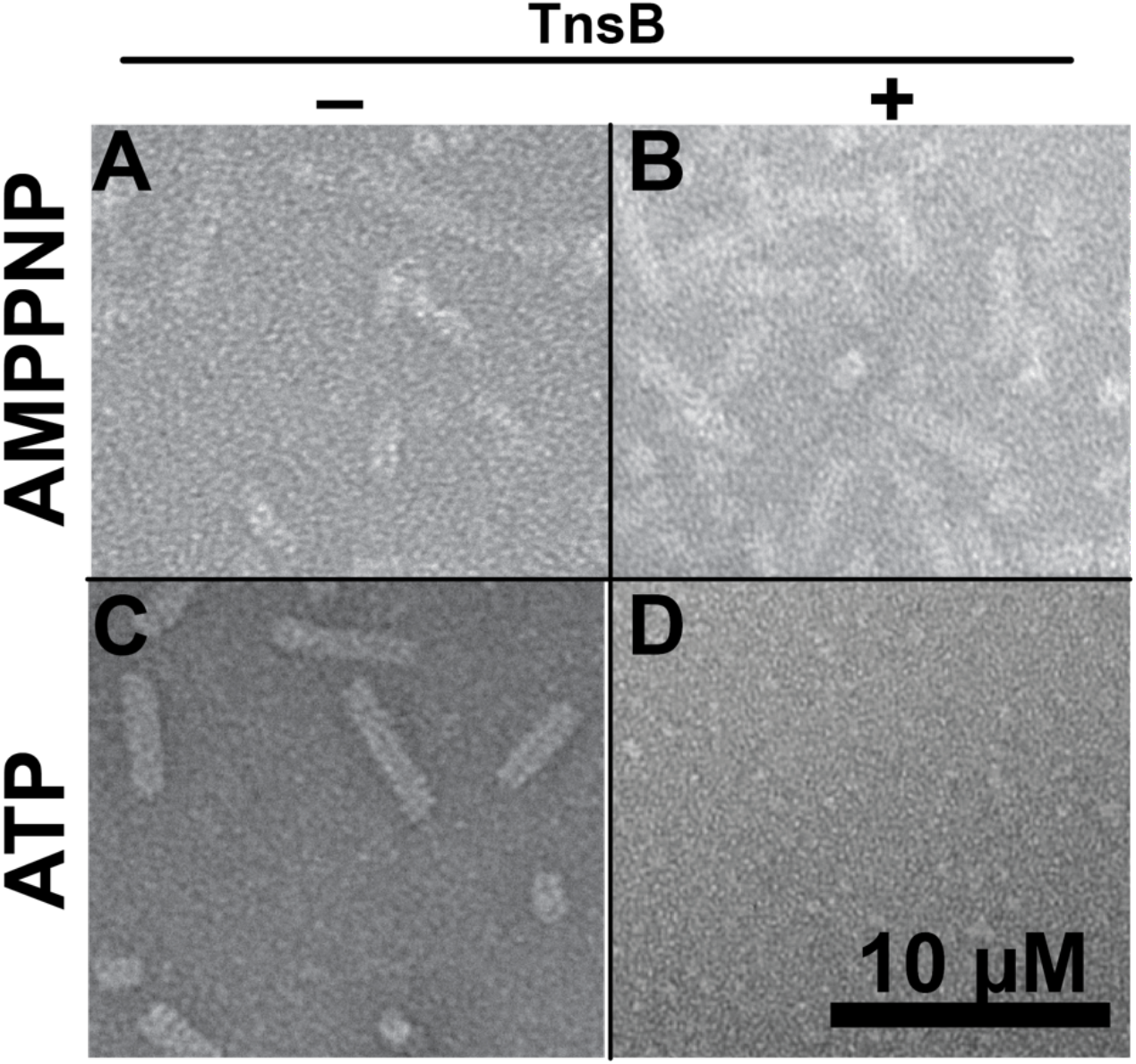
Filament assembly and disassembly of TnsC is coupled with ATP binding and TnsB-mediated ATP hydrolysis. Filament disassembly does not occur with a non-hydrolyzable analog. (**A-B**) Negative stain images of AMPPNP or (**C-D**) ATP-bound TnsC reveal that TnsC forms filaments in the presence of AMPPNP (**A**) and ATP (**C**) robustly. TnsB addition (right column) is sufficient to stimulate filament disassembly for ATP-TnsC (**D**) but not AMPPNP TnsC **(B)**

**Fig. S3.**
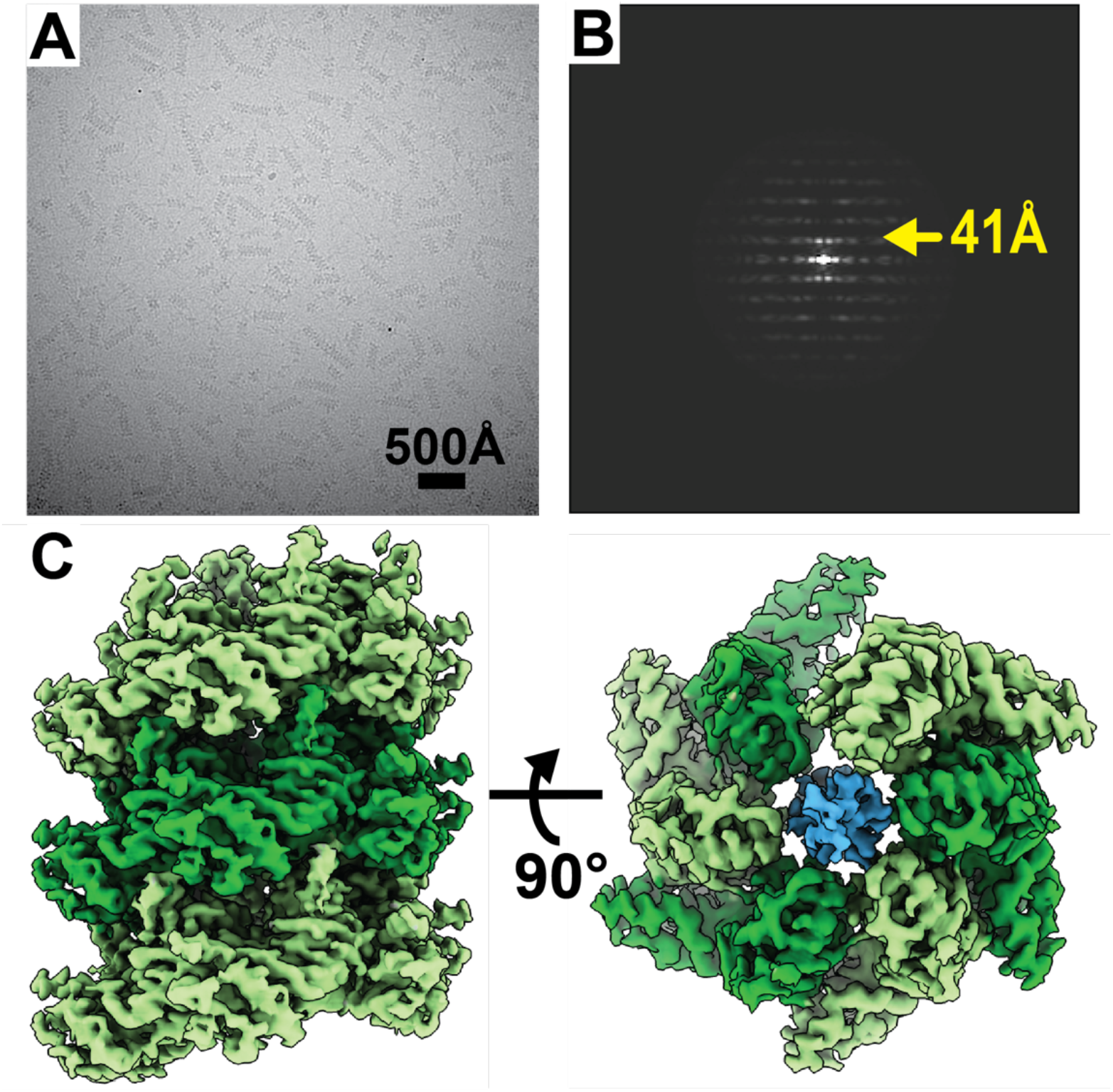
ATP-bound TnsC forms helical filaments that are identical to the ATPγS cryo-EM reconstruction. **A** Cryo-EM image of ATP-bound TnsC reveals helical filaments of variable length. **B.** Layer-line analysis of ATP-bound TnsC reveals a layer-line spacing of 41 Å. **C**. Cryo-EM reconstruction of ATP-TnsC reveals a right-handed helical filament encircling DNA, identical to what we observe for the case of non-hydrolyzable analog, ATPγS.

**Fig S4.**
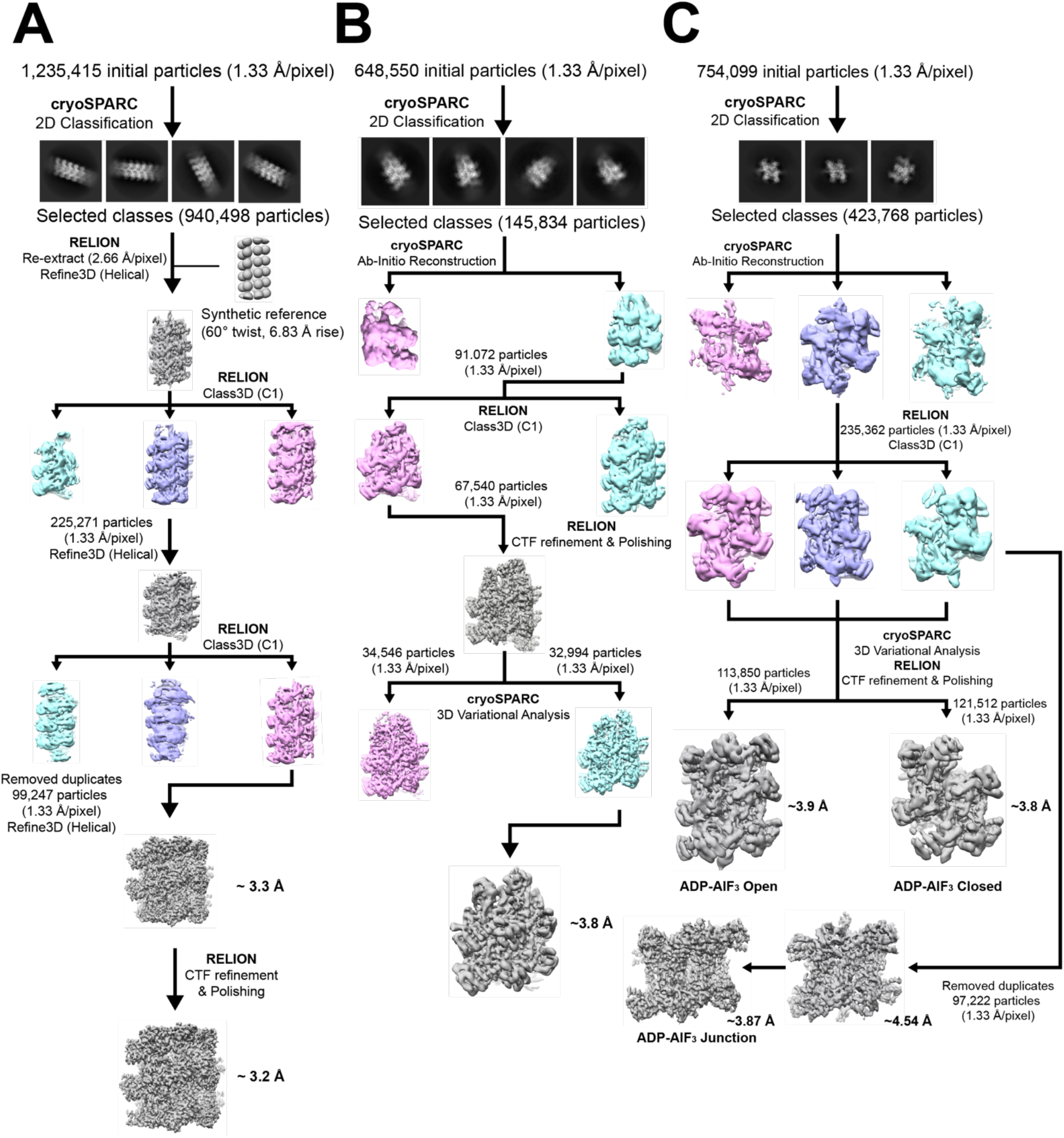
Cryo-EM Image processing workflow. **A.** ATPγS-bound TnsC filaments **B.** TniQ-bound TnsC filaments. **C.** ADP-AlF_3_ bound TnsC. For each workflow, 2D class averages are shown (top) followed by the RELION processing workflow, including 3D sorting, CTF-refinement, and particle polishing.

**Fig S5.**
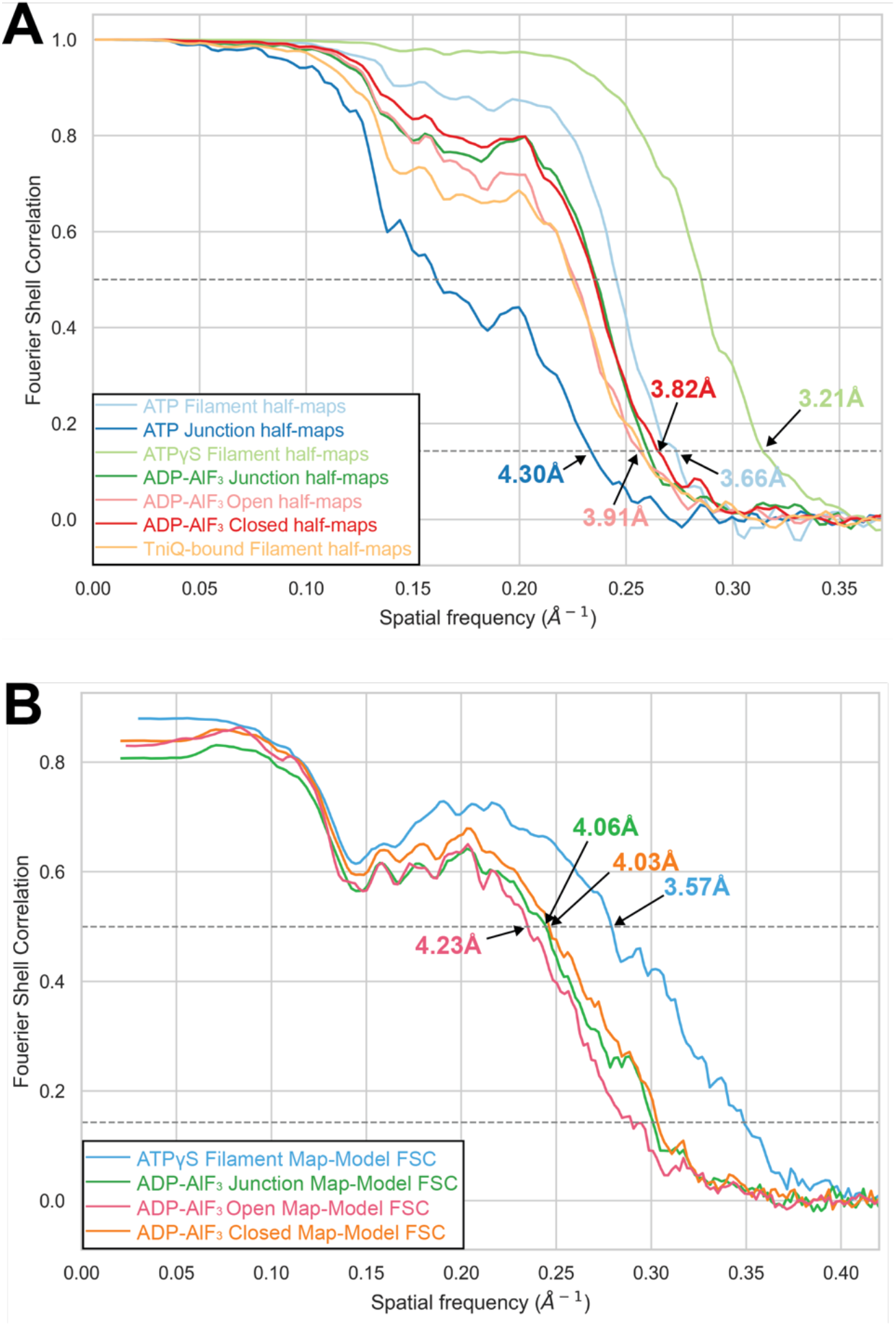
Fourier shell correlation from gold standard half-maps and model-map comparison. **A.** gold-standard FSC curve indicates all cryo-EM maps range between 3.2 – 4.3 Å resolution (0.143 cutoff). **B.** model-map FSC curves indicate that, based on the estimated resolution, the models fit the maps as expected given the resolution estimates (0.5 cutoff)

**Fig S6.**
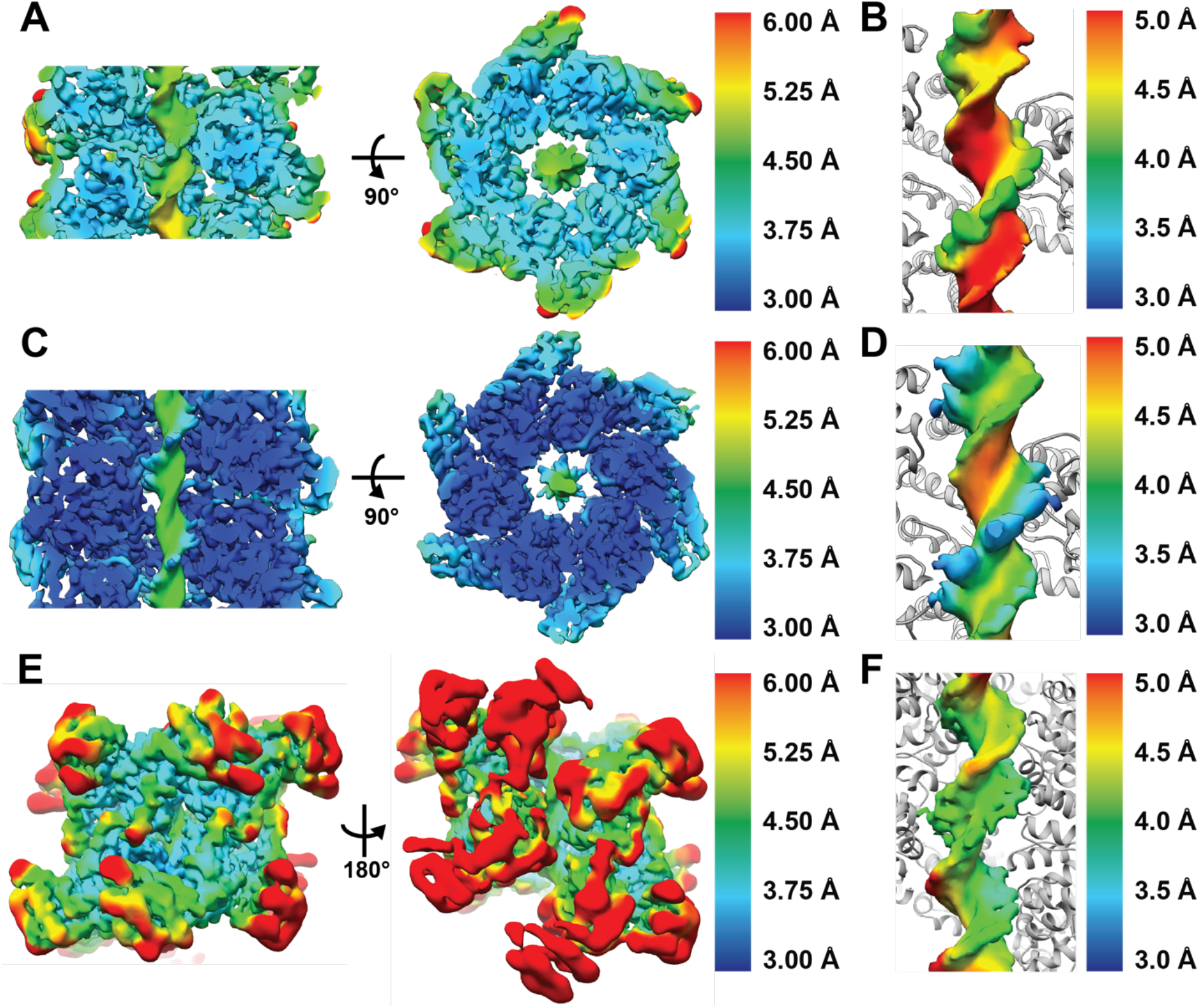
Local resolution of cryo-EM reconstructions of TnsC in various nucleotide-bound states: ATP, ATPγS and ADP·AlF_3_. **A.** ATP-bound filament. **B**. DNA of ATP-bound filament **C.** ATPγS bound filament. **D.** DNA from the ATPγS-bound filament. **E.** ADP·AlF_3_ cryo-EM reconstruction **F.** DNA from the ADP·AlF_3_ cryo-EM reconstruction. All maps are colored according to their local resolution, as estimated using Bsoft. Colors closer to blue indicate high-resolution (3 Å), whereas colors closer to red indicate low-resolution (5 Å+).

**Fig S7.**
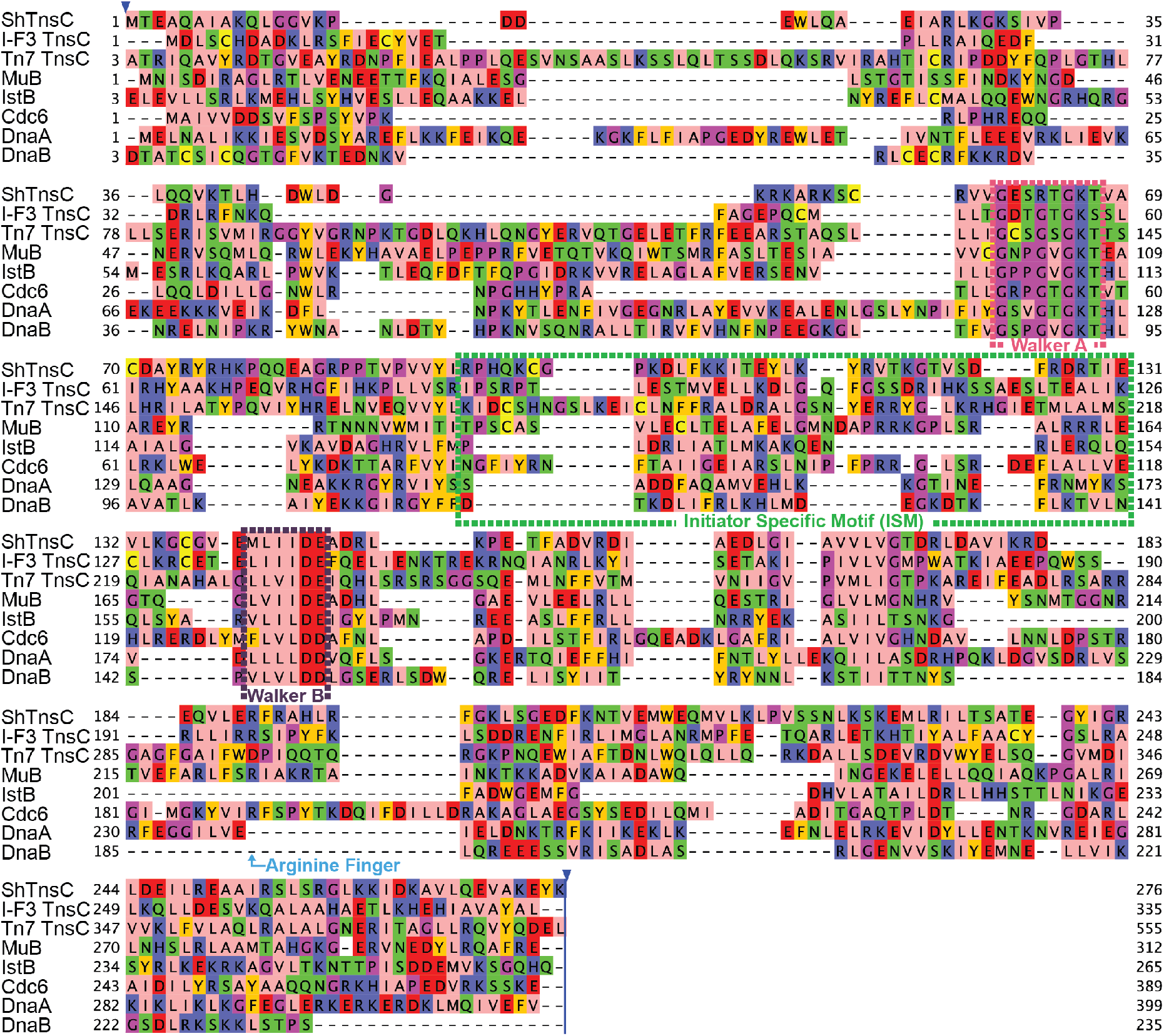
Multiple sequence alignment of TnsC homologs generated with MUSCLE, including: TnsC from ShCAST (referred to as ShTnsC), TnsC from Type IF-3b Tn7-CRISPR-Cas systems, prototypic Tn7 (labeled Tn7 TnsC), and related AAA+ proteins: MuB, IstB, Cdc6, DnaA, and DnaC.

**Fig S8.**
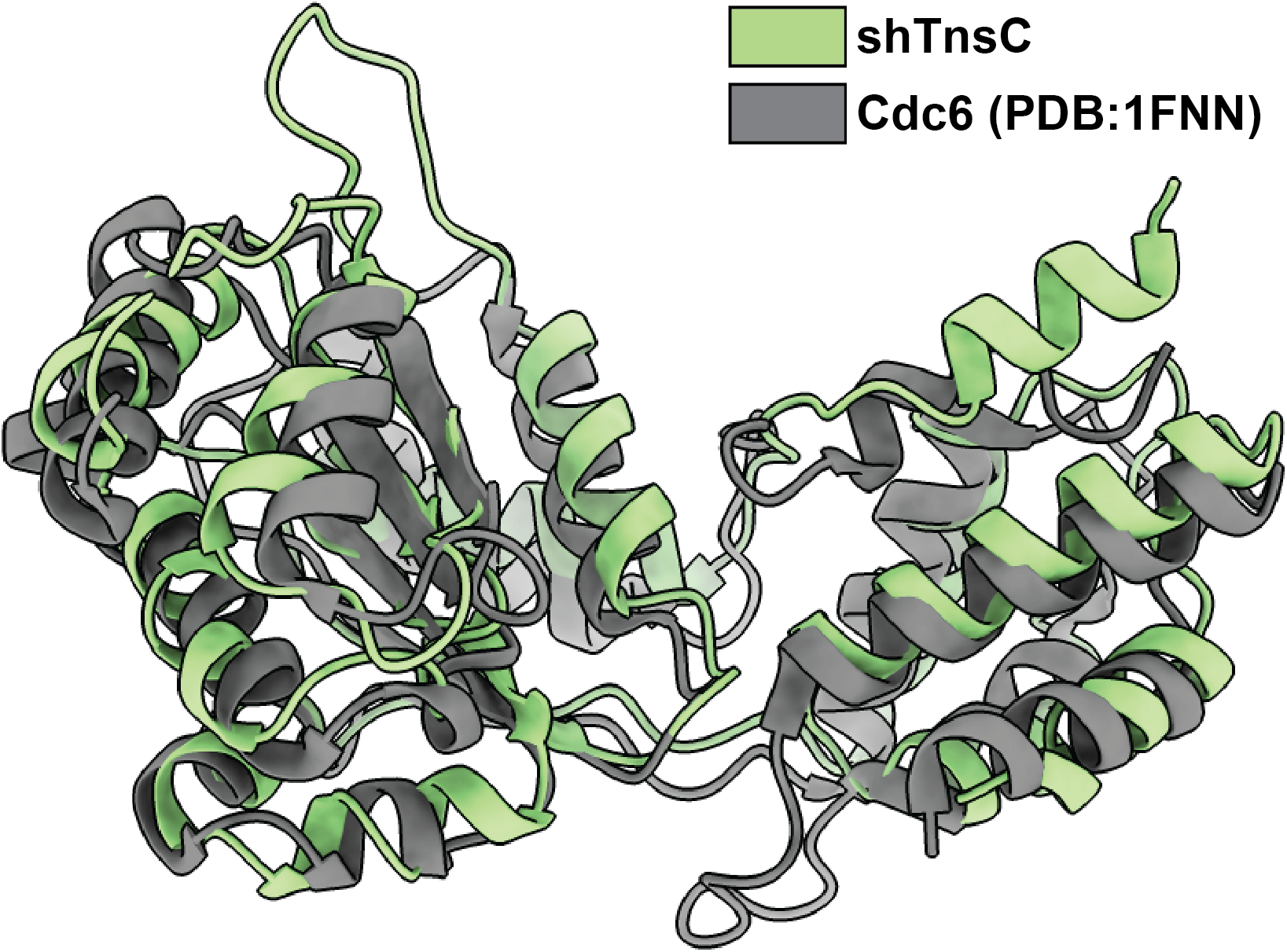
TnsC exhibits significant structural similarities to Cdc6. Green structure corresponds to ShCAST TnsC and gray model corresponds to the AAA+ domain of Cdc6 structure (PDB: 1FNN, residue 1 – 274). Global rmsd is 2.17 Å

**Fig S9.**
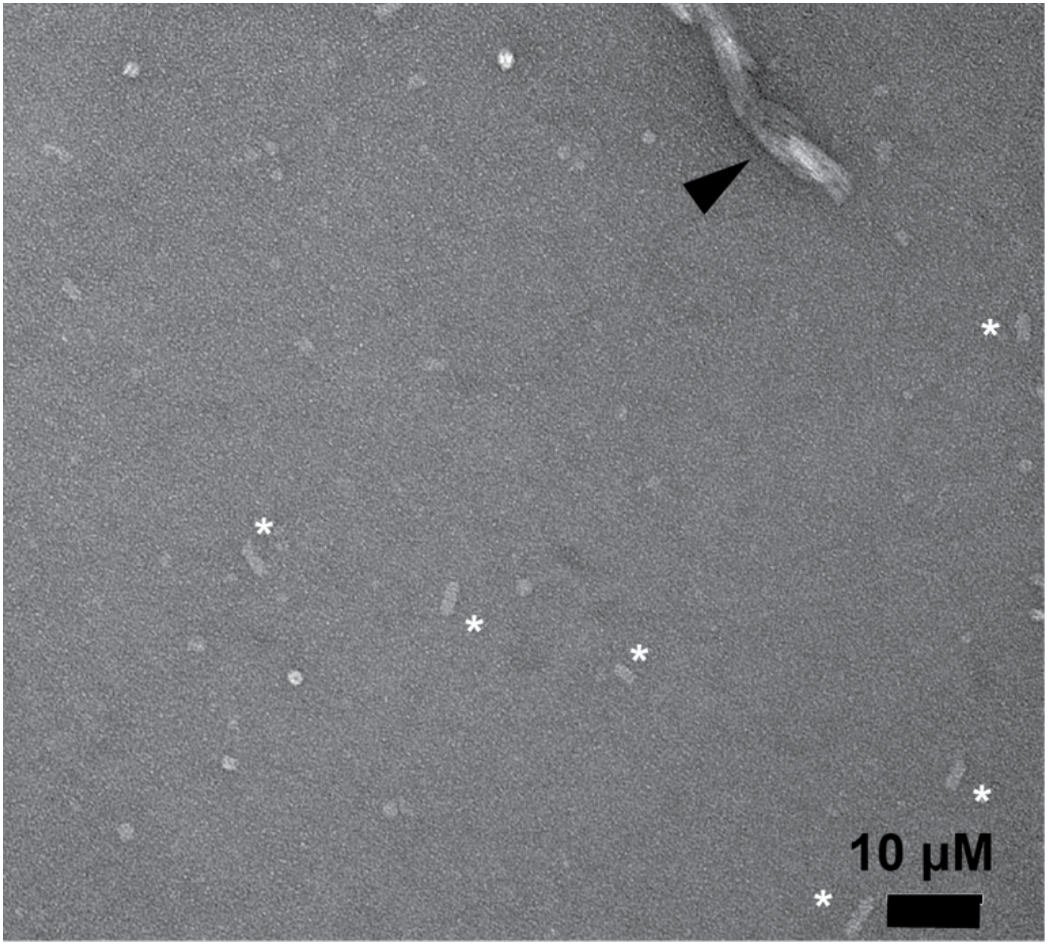
TnsC forms filaments in the presence of single-stranded DNA and ATP. Negative stain EM image reveals filaments in the presence of single-stranded DNA, albeit these filaments form less efficiently than in the case of double-stranded DNA. Black arrow indicates bundles of TnsC filaments. White asterisks indicate TnsC filaments.

**Fig S10.**
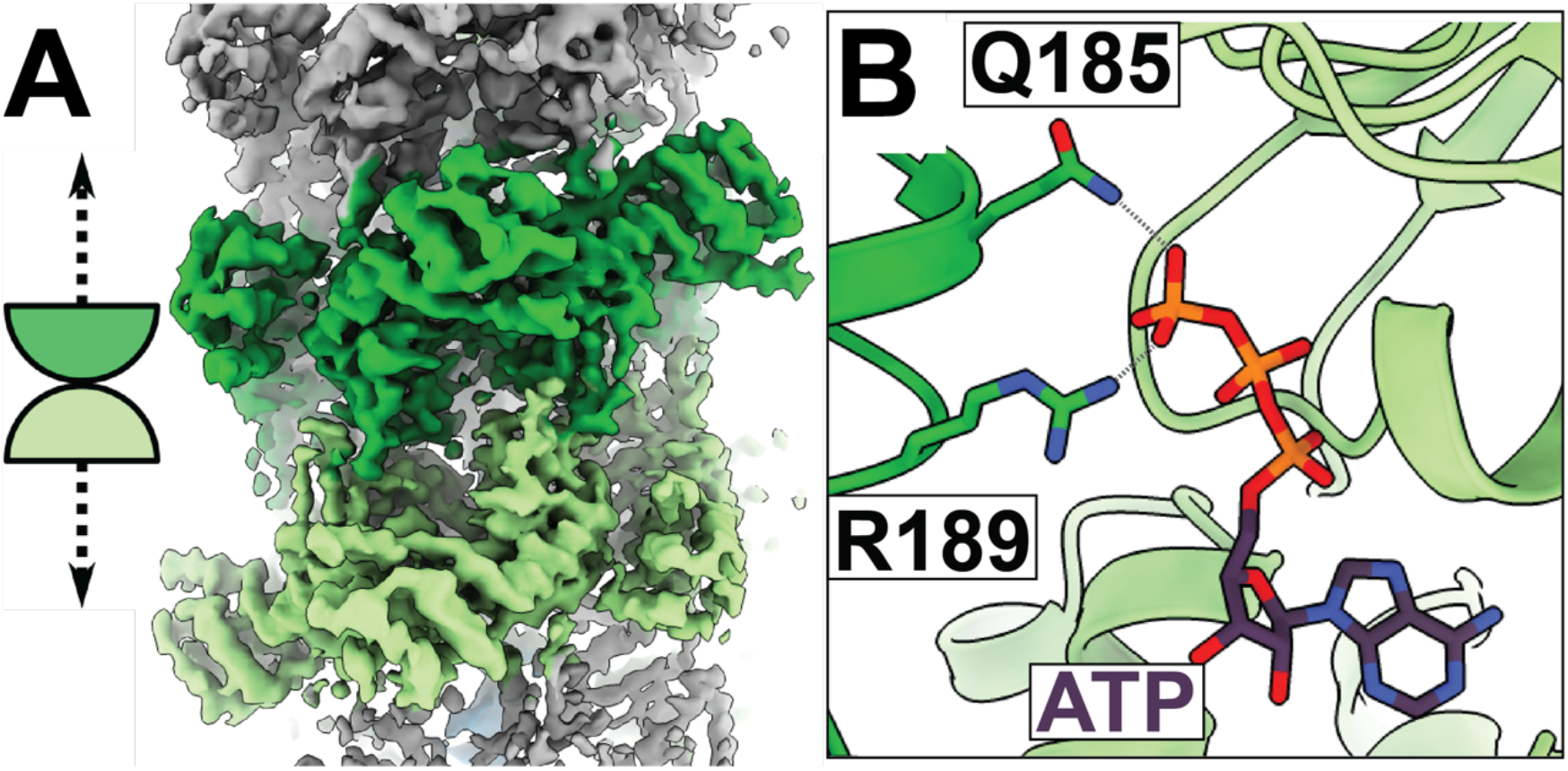
ATP structure of filaments colliding reveal a ‘head-to-head’ configuration. **A.** The green subunits highlight the junction, where two filaments collide, forming head-to-head interactions (i.e. the head face of one TnsC hexamer, colored dark green, is interacting with the head face of the opposing TnsC hexamer, colored light green). Additional gray density is observed on either side indicating that these junction structures occur in the middle of TnsC filaments. **B**. The ATP-binding site is consistent with what we observe in the ATPγS ATP-binding pocket, indicating that hydrolysis has not yet occurred.

**Fig S11.**
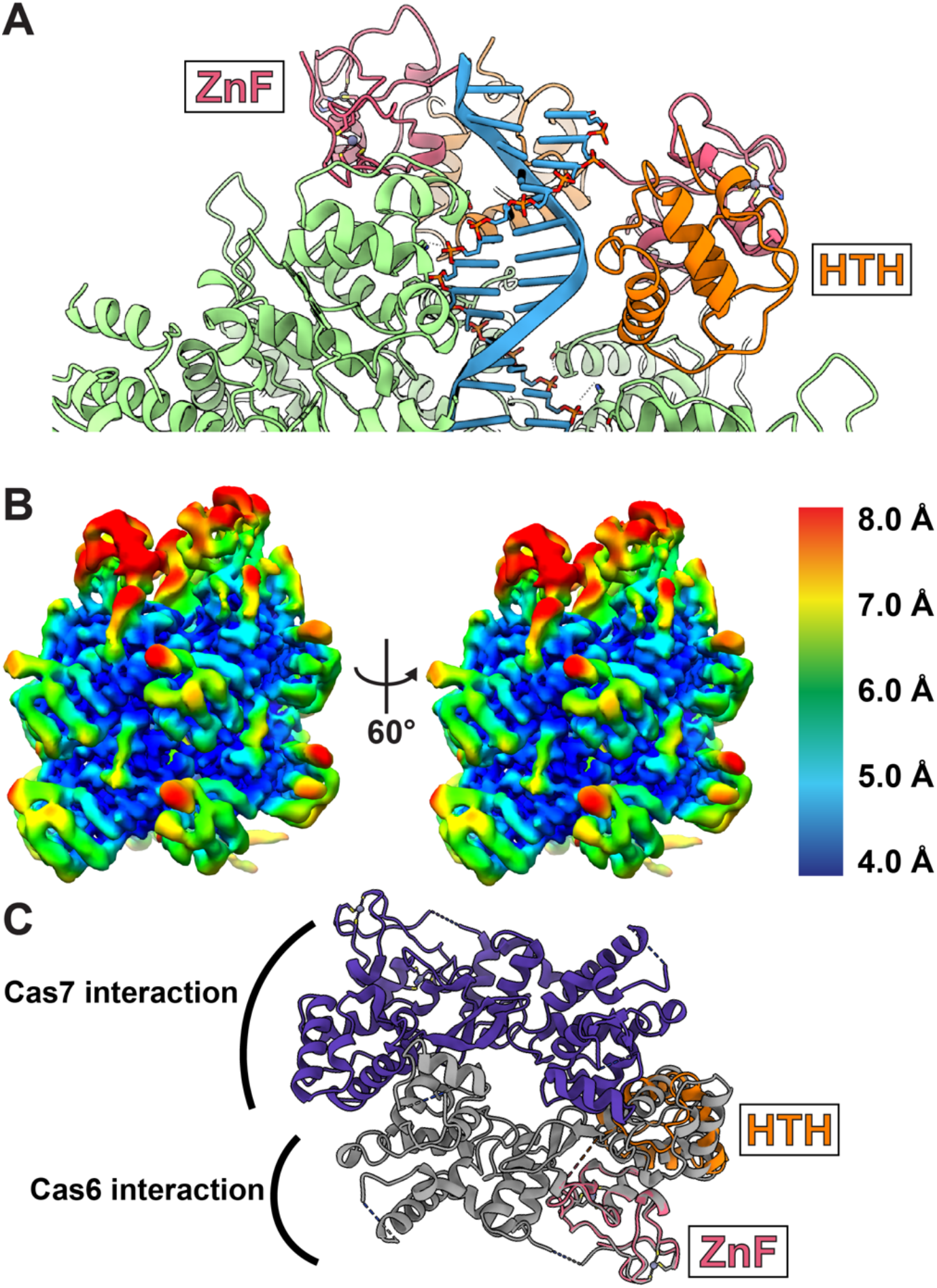
Characterization and structural analysis of the TniQ-TnsC cryo-EM map. **A.** TniQ (red and pink) mainly interacts with the strand of DNA (blue, ribbon representation) that is complementary to TnsC-interacting DNA-strand (stick representation). TnsC is shown in light green. **B.** Local resolution of TniQ ranges from 6-9 Å and is comparatively worse than the TnsC density (4-6 Å) in the TnsC-TniQ complex structure. Each voxel of the density map is colored according to the local resolution. **C**. The Type IF-3 TniQ is a homodimer (each subunit is colored purple or gray, PDB:6V9P). The TniQ dimer forms interactions with the Cascade complex (indicated). One Type IF-3 TniQ monomer (shown in gray) superimposes well with the modeled Type V TniQ monomer obtained from our cryo-EM reconstruction (orange and pink) (global rmsd is 2.19 Å).

**Fig S12.**
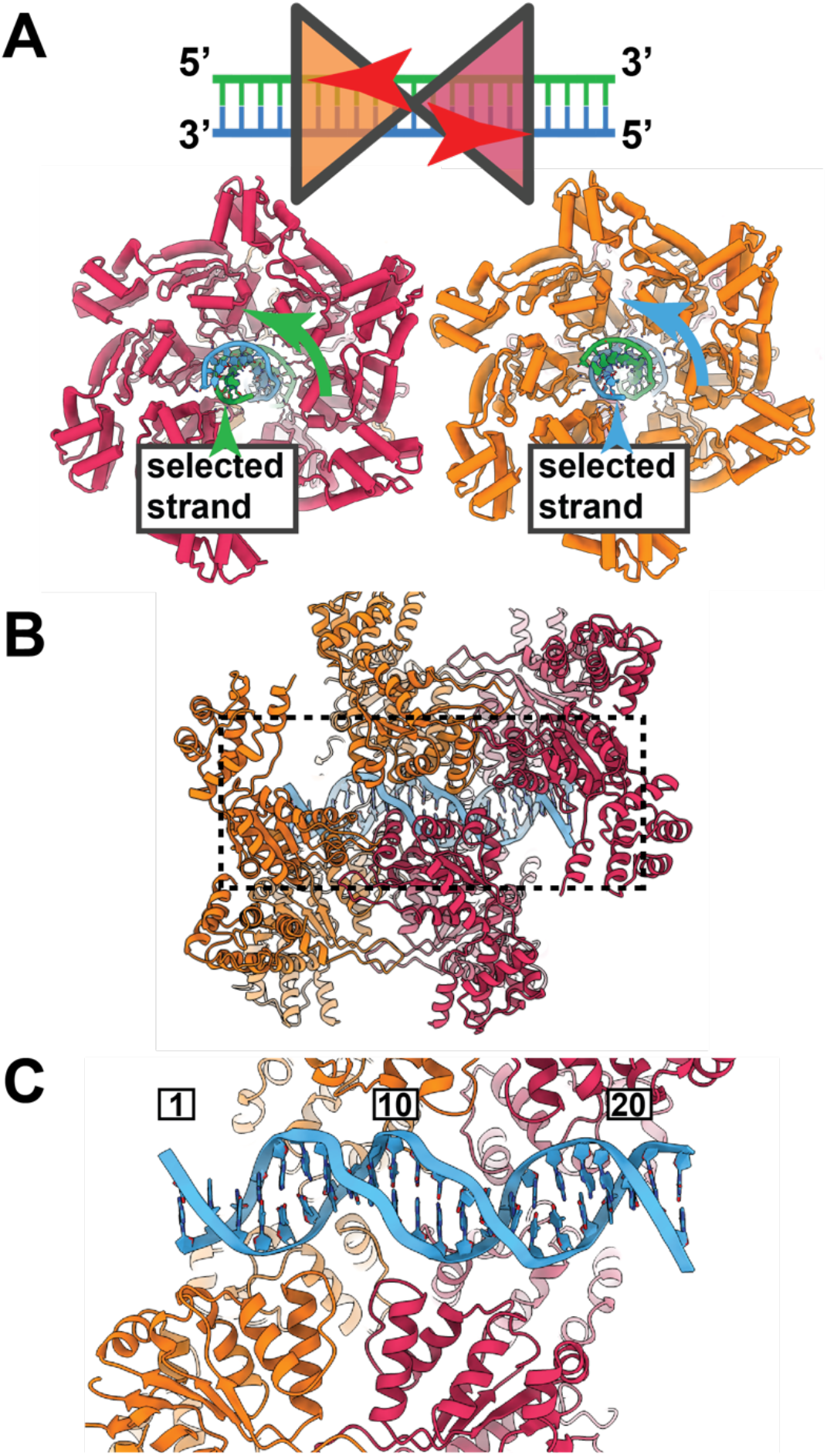
Each hexamer in the junction structure loads in a head-to-head orientation, resulting in binding of opposite DNA strands. **A.** Diagram of the hexamer orientation. Each hexamer is colored either pink or orange. Each strand of DNA is colored either green or blue. **B**. The head-to-head hexamer orientation forms an asymmetric complex which encircles and distorts DNA. **C.** The binding footprint spans 22 base-pairs. Binding of TnsC widens the major groove of DNA.

**Fig S13.**
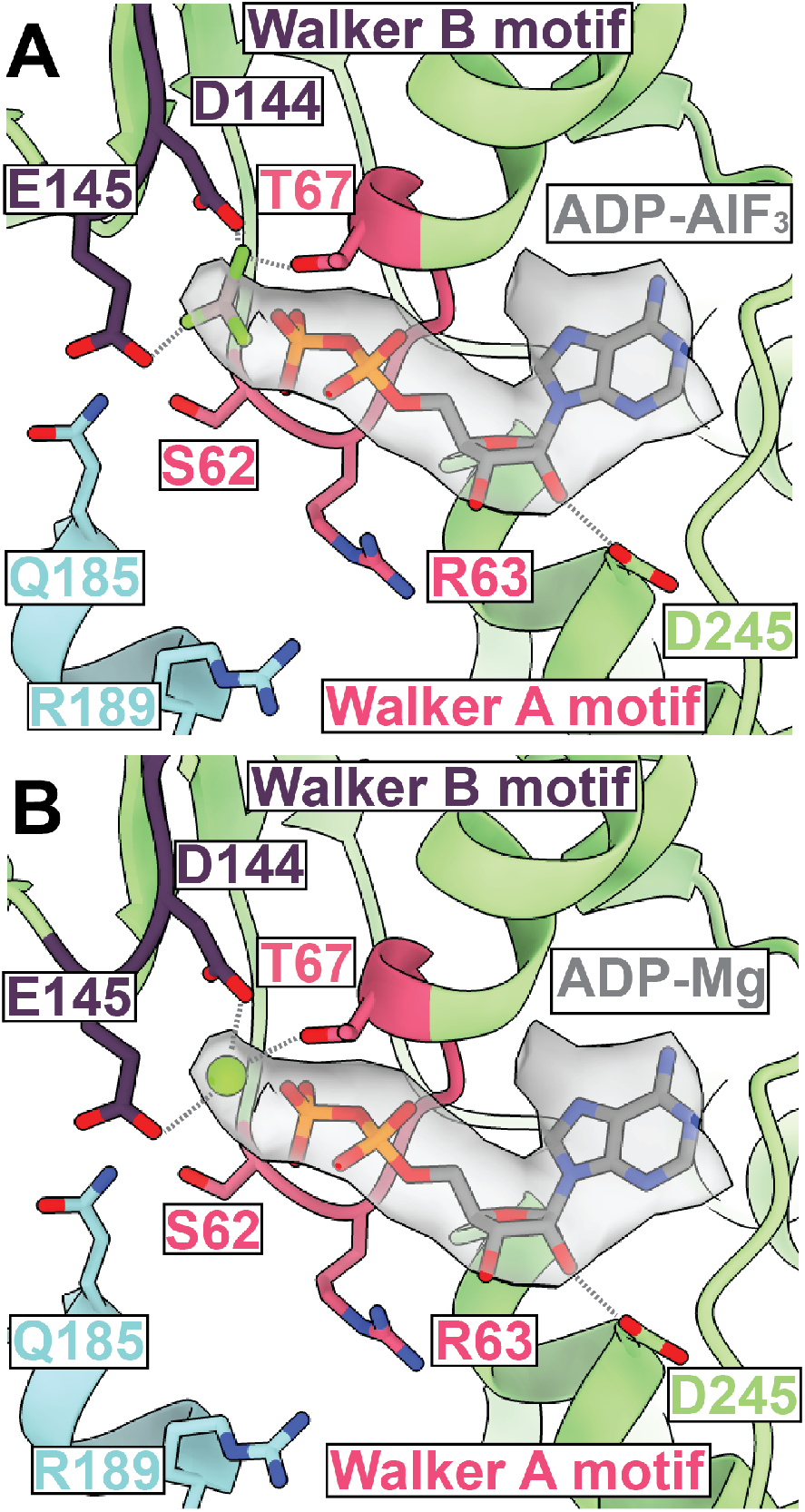
Nucleotide model docking of ATP binding pocket in ADP-AlF_3_ bound state of TnsC. **A.** ADP-AlF_3_ or **B.** ADP-Mg can both be modeled into the cryo-EM density corresponding to the nucleotide within the ATP-binding pocket.

**Table S1.** Summary of Cryo-EM datasets, including map and model statistics.

| | ATP TnsC<br>filament<br>EMD-23724 | ATP TnsC<br>junction<br>EMD-23725 | ATPγS TnsC<br>EMD-23720<br>PDB 7M99 | ADP•AIF<sub>3</sub><br>TnsC<br>EMD-23721<br>PDB 7M9A | ADP•AIF<sub>3</sub><br>open TnsC<br>EMD-23723<br>PDB 7M9C | ADP•AIF<sub>3</sub><br>closed TnsC<br>EMD-23722<br>PDB 7M9B | TniQ-TnsC<br>EMD-23726 |
| --- | --- | --- | --- | --- | --- | --- | --- |
| **Data Collection and Processing** |  |  |  |  |  |  |  |
| Magnification | 63,000 | 63,000 | 63,000 | 63,000 | 63,000 | 63,000 | 63,000 |
| Voltage (kV) | 200 | 200 | 200 | 200 | 200 | 200 | 200 |
| Electron<br>Exposure<br>(e<sup>-</sup> Å<sup>-2</sup>) | 45 | 45 | 40 | 45 | 45 | 45 | 50 |
| Defocus<br>range (μm) | -1 μm to<br>-2.5 μm | -1 μm to<br>-2.5 μm | -1 μm to<br>-2.5 μm | -1 μm to<br>-2.5 μm | -1 μm to<br>-2.5 μm | -1 μm to<br>-2.5 μm | -1 μm to<br>-2.5 μm |
| Pixel Size (Å) | 1.33 | 1.33 | 1.33 | 1.33 | 1.33 | 1.33 | 1.33 |
| Symmetry<br>imposed | helical | C1 | helical | C1 | C1 | C1 | C1 |
| Initial particle<br>images (no.) | 579,028 | 459,871 | 1,235,415 | 754,099 | 754,099 | 754,099 | 648,550 |
| Final particle<br>images (no.) | 33,312 | 57,363 | 99,247 | 97,222 | 113,850 | 121,512 | 34,546 |
| Map<br>resolution (Å)<br>/ FSC<br>threshold | 3.7 / 0.143 | 4.2 / 0.143 | 3.2 / 0.143 | 3.9 / 0.143 | 4.0 / 0.143 | 3.8 / 0.143 | 3.9 / 0.143 |
| Map<br>resolution<br>range (Å) | 3.5 – 6.0 | 4.0 – 8.5 | 3.0 – 4.0 | 3.5 – 6.5 | 3.5 – 6.5 | 3.5 – 6.5 | 3.5 – 7.5 |
| **Refinement** |  |  |  |  |  |  |  |
| Initial model<br>used | - | - | 6AZ0 | - | - | - | - |
| Model<br>resolution (Å)<br>/ FSC<br>threshold | N/A | N/A | 3.6 / 0.5 | 4.1 / 0.5 | 4.2 / 0.5 | 4.0 / 0.5 | N/A |
| Map<br>sharpening B<br>factor (Å<sup>2</sup>) | -41 | N/A | -50 | -51 | N/A | N/A | N/A |
| **Model composition** |  |  |  |  |  |  |  |
| Non-<br>hydrogen<br>atoms | - | - | 17,539 | 26,151 | 30,531 | 26,151 | - |
| Protein<br>residues | - | - | 2,056 | 3,084 | 3,598 | 3,084 | - |
| Ligands | - | - | 14 | 10 | 10 | 10 | - |
| **R.M.S. deviations** |  |  |  |  |  |  |  |
| Bond lengths<br>(Å) | - | - | 0.014 | 0.004 | 0.004 | 0.004 | - |
| Bond angles<br>(°) | - | - | 1.432 | 0.931 | 0.921 | 0.931 | - |
| **Validation** |  |  |  |  |  |  |  |
| MolProbity<br>Score | - | - | 1.45 | 1.54 | 1.62 | 1.55 | - |
| Clashscore | - | - | 8.27 | 7.28 | 9.34 | 7.57 | - |
| Poor rotamers (%) | - | - | 0.39 | 0.11 | 0.10 | 0.11 | - |
| **Ramachandran Plot** |  |  |  |  |  |  |  |
| Favored (%) | - | - | 98 | 97 | 97 | 97 | - |
| Allowed (%) | - | - | 2 | 3 | 3 | 3 | - |
| Disallowed (%) | - | - | 0 | 0 | 0 | 0 | - |

References (31–46)

### Movie S1

3D variability analysis reveals the conformational changes in the ADP-AlF3 sample. We are able to visualize the structural changes between the ‘closed’ and the ‘open’ conformation within the ADP-AlF3 ensemble.

